# Structural characterization of encapsulated ferritin provides insight into iron storage in bacterial nanocompartments

**DOI:** 10.1101/063495

**Authors:** Didi He, Sam Hughes, Sally Vanden-Hehir, Atanas Georgiev, Kirsten Altenbach, Emma Tarrant, C. Logan Mackay, Kevin J. Waldron, David J. Clarke, Jon Marles-Wright

**Affiliations:** Institute of Quantitative Biology, Biochemistry and Biotechnology, School of Biological Sciences, The University of Edinburgh, Max Born Crescent, Edinburgh, EH9 3BF; The School of Chemistry, The University of Edinburgh, Joseph Black Building, David Brewster Road, Edinburgh, Scotland. EH9 3FJ; Institute for Cell and Molecular Biosciences, Newcastle University, Newcastle upon Tyne, NE2 4HH; School of Biology, Newcastle University. Newcastle upon Tyne, NE1 7RU

## Abstract

Ferritins are ubiquitous proteins that oxidise and store iron within a protein shell to protect cells from oxidative damage. We have characterized the structure and function of a new member of the ferritin superfamily that is sequestered within an encapsulin capsid. We show that this encapsulated ferritin (EncFtn) has two main alpha helices, which assemble in a metal dependent manner to form a ferroxidase centre at a dimer interface. EncFtn adopts an open decameric structure that is topologically distinct from other ferritins. While EncFtn acts as a ferroxidase, it cannot mineralize iron. Conversely, the encapsulin shell associates with iron, but is not enzymatically active, and we demonstrate that EncFtn must be housed within the encapsulin for iron storage. This encapsulin nanocompartment is widely distributed in bacteria and archaea and represents a distinct class of iron storage system where the oxidation and mineralization of iron are distributed between two proteins.

## Introduction

Encapsulin nanocompartments are a family of proteinaceous metabolic compartments that are widely distributed in bacteria and archaea^1–4^. They share a common architecture, comprising an icosahedral shell formed by the oligomeric assembly of a protein, encapsulin, that is structurally related to the HK97 bacteriophage capsid protein gp5^5^. Gp5 is known to assemble as a 66 nm diameter icosahedral shell of 420 subunits. In contrast, both the *Pyrococcus furiosus*^2^ and *Myxococcus xanthus*^3^ encapsulin shell-proteins form 32 nm icosahedra with 180 subunits; while the *Thermotoga maritima*^6^ encapsulin is smaller still with a 25 nm, 60-subunit icosahedron. The high structural similarity of the encapsulin shell-proteins to gp5 suggests a common evolutionary origin for these proteins^3^.

The genes encoding encapsulin proteins are found downstream of genes for dye-dependent peroxidase (DyP) family enzymes^7^, or encapsulin-associated ferritins (EncFtn)^8^. Enzymes in the DyP family are active against polyphenolic compounds such as azo dyes and lignin breakdown products; although their physiological function and natural substrates are not known^7^. Ferritin family proteins are found in all kingdoms and have a wide range of activities, including ribonucleotide reductase^9^, protecting DNA from oxidative damage^10^, and iron storage^11^. The classical iron storage ferritin nanocages are essential in eukaryotes and play a central role in iron homeostasis, where they protect the cell from toxic free Fe^2+^ by oxidizing it and storing the resulting Fe^3+^ as ferrihydrite minerals within their central cavity.

The encapsulin-associated enzymes are sequestered within the icosahedral shell through interactions between the shell's inner surface and a short localization sequence (Gly-Ser-Leu-Lys) appended to their C-termini^1^. This motif is well-conserved, and addition of this sequence to heterologous proteins is sufficient to direct them to the interior of encapsulins^12,13^.

A recent study of the *Myxococcus xanthus* encapsulin showed that it sequesters a number of different EncFtn proteins and acts as an ‘iron-megastore’ to protect these bacteria from oxidative stress^14^. At 32 nm in diameter, it is much larger than other members of the ferritin superfamily, such as the 12 nm 24-subunit classical ferritin nanocage and the 8 nm 12-subunit Dps complex^10,15^; and is thus capable of sequestering up to ten times more iron than these ferritins^3^. The primary sequences of EncFtn proteins have Glu-X-X-His metal coordination sites, which are shared features of the ferritin family proteins^15^. Secondary structure prediction identifies two major a-helical regions in these proteins; this is in contrast to other members of the ferritin superfamily, which have four major α-helices (Supplementary File 1). The ‘half-ferritin’ primary sequence of the EncFtn family and their association with encapsulin nanocompartments suggests a distinct biochemical and structural organization to other ferritin family proteins. The *R. rubrum* EncFtn protein (Rru_A0973) shares 33% protein sequence identity with the *M. Xanthus* (MXAN_4464), 53% with the *T. maritima* (Tmari_0787), and 29% with the *P. furiosus* (PF1192) homologues. The GXXH motifs are strictly conserved in each of these species (Supplementary file 1).

Here we investigate the structure and biochemistry of EncFtn in order to understand iron storage within the encapsulin nanocompartment. We have produced recombinant encapsulin (Enc) and EncFtn from the aquatic purple-sulfur bacterium *Rhodospirillum rubrum*, which serves as a model organism for the study of the control of the bacterial nitrogen fixation machinery^16^, in *Escherichia coli*. Analysis by transmission electron microscopy (TEM) indicates that their coexpression leads to the production of an icosahedral nanocompartment with encapsulated EncFtn. The crystal structure of a truncated hexahistidine-tagged variant of the EncFtn protein (EncFtn_sH_) shows that it forms a decameric structure with an annular ‘ring-doughnut’ topology, which is distinct from the four-helical bundles of the 24meric ferritins^17^ and dodecahedral DPS proteins^10^. We identify a symmetrical iron bound ferroxidase center (FOC) formed between subunits in the decamer and additional metal-binding sites close to the center of the ring and on the outer surface. We also demonstrate the metal-dependent assembly of EncFtn decamers using native PAGE, analytical gel-filtration, and native mass spectrometry. Biochemical assays show that EncFtn is active as a ferroxidase enzyme. Through site-directed mutagenesis we show that the conserved glutamic acid and histidine residues in the FOC influence protein assembly and activity. We use our combined structural and biochemical data to propose a model for the EncFtn-catalyzed sequestration of iron within the encapsulin shell.

## Results

### Assembly of *Rhodospirillum rubrum* EncFtn encapsulin nanocompartments in *E. coli*

We produced recombinant *Rhodospirillum rubrum* encapsulin nanocompartments in *E. coli* by coexpression of the encapsulin (Rru_A0974) and EncFtn (Rru_A0973) proteins, and purified these by sucrose gradient ultra-centrifugation (Figure 1A)^1^. TEM imaging of uranyl acetate-stained samples revealed that, when expressed in isolation, the encapsulin protein forms empty compartments with an average diameter of 24 nm (Figure 1B and Figure 1-figure supplement 1A/C), consistent with the appearance and size of the *T. maritima* encapsulin^1^. We were not able to resolve any higher-order structures of EncFtn by TEM. Protein purified from co-expression of the encapsulin and EncFtn resulted in 24 nm compartments with regions in the center that exclude stain, consistent with the presence of the EncFtn within the encapsulin shell (Figure 1C and Figure 1-figure supplement 1B/C).

### *R. rubrum* EncFtn forms a metal-ion stabilized decamer in solution

We purified recombinant *R. rubrum* EncFtn as both the full-length sequence (140 amino acids) and a truncated C-terminal hexahistidine-tagged variant (amino acids 1-96 plus the tag; herein EncFtn_sH_). In both cases the elution profile from size-exclusion chromatography (SEC) displayed two peaks (Figure 2A). SDS-PAGE analysis of fractions from these peaks showed that the high molecular weight peak was partially resistant to SDS and heat-induced denaturation; in contrast, the low molecular weight peak was consistent with monomeric mass of 13 kDa (Figure 2B). MALDI peptide mass fingerprinting of these bands confirmed the identity of both as EncFtn. Inductively coupled plasma mass spectrometry (ICP-MS) analysis of the SEC fractions showed 100 times more iron in the oligomeric fraction than the monomer (Figure 2A, blue scatter points; Table 1), suggesting that EncFtn oligomerization is associated with iron binding. In order to determine the iron-loading stoichiometry in the EncFtn complex, further ICP-MS experiments were performed using selenomethionine-labelled protein EncFtn (Table 1). In these experiments we observed sub-stoichiometric metal binding, which is in contrast to the classical ferritins^18^. Size exclusion chromatography with multi-angle laser light scattering (SEC-MALLS) analysis of samples taken from each peak gave calculated molecular weights consistent with a decamer for the high molecular weight peak and a monomer for the low molecular weight peak (Figure 2C, Table 2).

We purified EncFtn_sH_ from *E. coli* grown in minimal media with or without the addition of 1 mM Fe(NH_4_)_2_(SO_4_)_2_. The decamer to monomer ratio in the sample purified from cells grown in iron-supplemented media was 4.5, while that from the iron-free media was 0.11, suggesting that iron induces the oligomerization of EncFtn_sH_ *in vivo* (Figure 3A, Table 3). To test the metal-dependent oligomerization of EncFtn_sH_ *in vitro*, we incubated the protein with various metal cations and subjected samples to analytical SEC and non-denaturing PAGE. Of the metals tested, only Fe^2+^, Zn^2+^ and Co^2+^ induced the formation of significant amounts of the decamer (Figure 3B, Figure 3-supplement 1/2). While Fe^2+^ induces multimerization of EncFtn_sH_, Fe^3+^ in the form of FeCl_3_ does not have this effect on the protein, highlighting the apparent preference this protein has for the ferrous form of iron. To determine if the oligomerization of EncFtn_sH_ was concentration dependent we performed analytical SEC at 90 and 700 μM protein concentration (Figure 3C). At the higher concentration, no increase in the decameric form of EncFtn was observed; however, the shift in the major peak from the position of the monomer species indicated a tendency to dimerize at high concentration.

### Crystal structure of EncFtn_sH_

We determined the crystal structure of EncFtn_sH_ by molecular replacement to 2.0 Å resolution (see Table 1 for X-ray data collection and refinement statistics). The crystallographic asymmetric unit contained thirty monomers of EncFtn with visible electron density for residues 7 – 96 in each chain. The protein chains were arranged as three identical annular decamers, each with D5 symmetry. The decamer has a diameter of 7 nm and thickness of 4 nm (Figure 4A). The monomer of EncFtn has an N-terminal 3_10_-helix that precedes two 4 nm long antiparallel α-helices arranged with their long axes at 25° to each other; these helices are followed by a shorter 1.4 nm helix projecting at 70° from α2 (Figure 4B). The C-terminal region of the crystallized construct extends from the outer circumference of the ring, indicating that the encapsulin localization sequence in the full-length protein is on the exterior of the ring and is thus free to interact with its binding site on the encapsulin shell protein^1^.

The monomer of EncFtn_sH_ forms two distinct dimer interfaces within the decamer (Figure 4 C/D). The first dimer is formed from two monomers arranged antiparallel to each other, with a1 from each monomer interacting along their lengths and α3 interdigitating with α2 and α3 of the partner chain. This interface buries one third of the surface area from each partner and is stabilized by thirty hydrogen bonds and fourteen salt bridges (Figure 4C). The second dimer interface forms an antiparallel four-helix bundle between helices 1 and 2 from each monomer (Figure 4C). This interface is less extensive than the first and is stabilized by twenty-one hydrogen bonds, six salt bridges, and a number of metal ions.

The arrangement of ten monomers in alternating orientation forms the decamer of EncFtn, which assembles as a pentamer of dimers (Figure 4A). Each monomer lies at 45° relative to the vertical central-axis of the ring, with the N-termini of alternating subunits capping the center of the ring at each end, while the C-termini are arranged around the circumference. The central hole in the ring is 2.5 nm at its widest in the center of the complex, and 1.5 nm at its narrowest point near the outer surface, although it should be noted that a number of residues at the N-terminus are not visible in the crystallographic electron density and these may occupy the central channel. The surface of the decamer has distinct negatively charged patches, both within the central hole and on the outer circumference, which form spokes through the radius of the complex (Figure 4-figure supplement 1).

### EncFtn ferroxidase center

The electron density maps of the initial EncFtn_sH_ model displayed significant positive peaks in the mFo-DFc map at the center of the 4-helix bundle dimer (Figure 5-figure supplement 1). Informed by the ICP-MS data indicating the presence of iron in the protein we collected diffraction data at the experimentally determined iron absorption edge (1.739 Å) and calculated an anomalous difference Fourier map using this data. Inspection of this map showed two 10-sigma peaks between residues Glu32, Glu62 and His65 of two adjacent chains, and a statistically smaller 5-sigma peak between residues Glu31 and Glu34 of the two chains. Modeling divalent metal ions into these peaks and refinement of the anomalous scattering parameters allowed us to identify these as two iron ions and a calcium ion respectively (Figure 5A). An additional region of asymmetric electron density near the di-iron binding site in the mFo-DFc map was modeled as glycolic acid, a breakdown product of the PEG 3350 used for crystallization. This di-iron center has an Fe-Fe distance of 3.5 Å Fe-Glu-O distances between 2.3 and 2.5 Å, and Fe-His-N distances of 2.5 Å (Figure 5B). This coordination geometry is consistent with the di-nuclear ferroxidase center (FOC) found in ferritin^19^. It is interesting to note that although we did not add any additional iron to the crystallization trials, the FOC was fully occupied with iron in the final structure, implying that this site has a very high affinity for iron.

The calcium ion coordinated by Glu31 and Glu34 adopts heptacoordinate geometry, with coordination distances of 2.5 Å between the metal ion and carboxylate oxygens of Glu31 and Glu34 (E31/34-site). A number of ordered solvent molecules are also coordinated to this metal ion at a distance of 2.5 Å. This heptacoordinate geometry is common in crystal structures with calcium ions (Figure 5C)^20^. While ICP-MS indicated that there were negligible amounts of calcium in the purified protein, the presence of 140 mM calcium acetate in the crystallization mother liquor favors the coordination of calcium at this site. The fact that the protein does not multimerize in solution in the presence of Fe^3+^ may indicate that these metal binding sites have a lower affinity for the ferric form of iron, which is the product of the ferroxidase reaction. A number of additional low-occupancy metal-ions were present at the outer circumference of at least one decamer in the asymmetric unit (Fig. 5D). These ions are coordinated by His57, Glu61 and Glu64 from both chains in the FOC dimer and are 4.5 Å apart; Fe-Glu-O distances are between 2.5 and 3.5 Å and the Fe-His-N distances are 4 and 4.5 Å.

Structural alignment of the di-iron binding site of EncFtn_sH_ to the FOC of *Pseudo-nitzschia multiseries* ferritin (PmFtn, PDB ID: 4ITW) reveals a striking similarity between the metal binding sites of EncFtn_sH_ and the classical ferritins (Fig. 6A). The di-iron site of EncFtn_sH_ is by necessity symmetrical, as it is formed through a dimer interface, while the FOC of ferritin does not have these constraints and varies in different species at a position equivalent to His65 of the second EncFtn monomer in the FOC interface (H65’) (Figure 6A). Structural superimposition of the FOCs of ferritin and EncFtn brings the four-helix bundle of the ferritin fold into close alignment with the EncFtn dimer, showing that the two families of proteins have essentially the same architecture around the di-iron center (Figure 6B). The linker connecting helices 2 and 3 of ferritin is congruent with the start of the C-terminal helix of one EncFtn monomer and the N-terminal 3_10_ helix of the second monomer (Figure 6C).

### Mass spectrometry of the EncFtn assembly

In order to confirm the assignment of the oligomeric state of EncFtn_sH_ and investigate further the Fe^2+^-dependent assembly, we used native nanoelectrospray ionization (nESI) and ion-mobility mass spectrometry (IM-MS). As described above, by recombinant production of EncFtn_sH_ in minimal media we were able to limit the bioavailability of iron. Native MS analysis of EncFtn_sH_ produced in this way displayed a charge state distribution consistent with an EncFtn_sH_ monomer (blue circles, Figure 7A1) with an average neutral mass of 13,194 Da, in agreement with the predicted mass of the EncFtn_sH_ protein (13,194.53 Da). Under these conditions, no significant higher order assembly was observed and the protein did not have any coordinated metal ions. Titration with Fe^2+^ directly before native MS analysis resulted in the appearance of a new charge state distribution, consistent with an EncFtn_sH_ decameric assembly (+22 to +26; 132.65 kDa) (Figure 7A2/3). After instrument optimization, the mass resolving power achieved was sufficient to assign iron-loading in the complex to between 10 and 15 Fe ions per decamer (Figure 7B, inset top right), consistent with the presence of 10 irons in the FOC and the coordination of iron in the E31/34-site occupied by calcium in the crystal structure (*Δ*mass ~0.67 kDa). MS analysis of EncFtn after addition of further Fe^2+^ did not result in iron loading above this stoichiometry. Therefore, the extent of iron binding seen is limited to the FOC and E31/34 secondary metal binding site. These data suggest that the decameric assembly of EncFtn does not accrue iron in the same manner as classical ferritin, which is able to sequester more than 4000 iron ions within its nanocage^21^. Ion mobility analysis of the EncFtn_sH_ decameric assembly, collected with minimal collisional activation, suggested that it consists of a single conformation with a collision cross section (CCS) of 58.2 nm^2^ (Figure 7B). This observation is in agreement with the calculated CCS of 58.7 nm^2^ derived from our crystal structure of the EncFtn_sH_ decamer^22^. By contrast, IM-MS measurements of the monomeric EncFtn_sH_ at pH 8.0 under the same instrumental conditions revealed that the metal-free protein monomer exists in a wide range of charge states (+6 to +16) and adopts many conformations in the gas phase with collision cross sections ranging from 12 nm^2^ to 26 nm^2^ (Figure 7-figure supplement 1). These observations are indicative of an unstructured protein with little secondary or tertiary structure^23^. Thus, IM-MS studies highlight that higher order structure in EncFtn_sH_ is mediated/stabilized by metal binding, an observation that is in agreement with our solution studies. Taken together, these results suggest that di-iron binding, forming the FOC in EncFtn_sH_, is required to stabilize the 4-helix bundle dimer interface, essentially reconstructing the classical ferritin-like fold; once stabilized, these dimers readily associate as pentamers, and the overall assembly adopts the decameric ring arrangement observed in the crystal structure.

We subsequently performed gas phase disassembly of the decameric EncFtn_sH_ using collision-induced dissociation (CID) tandem mass spectrometry. Under the correct CID conditions, protein assemblies can dissociate with retention of subunit and ligand interactions, and thus provide structurally-informative evidence as to the topology of the original assembly; this has been termed ‘atypical’ dissociation^24^. For EncFtn_sH_, this atypical dissociation pathway was clearly evident; CID of the EncFtn_sH_ decamer resulted in the appearance of a dimeric EncFtn_sH_ subcomplex containing 0, 1, or 2 iron ions (Figure 7-figure supplement 2). In light of the crystal structure, this observation can be rationalized as dissociation of the EncFtn_sH_ decamer by disruption of the non-FOC interface with at least partial retention of the FOC interface and the FOC-Fe. Thus, this observation supports our crystallographic assignment of the overall topology of the EncFtn_sH_ assembly as a pentameric assembly of dimers with two iron ions located at the FOC dimer interface. In addition, this analysis provides evidence that the overall architecture of the complex is consistent in the crystal, solution and gas phases.

### Ferroxidase activity

In light of the identification of an iron-loaded FOC in the crystal structure of EncFtn and our native mass spectrometry data, we performed ferroxidase and peroxidase assays to demonstrate the catalytic activity of this protein. In addition, we also assayed equine apoferritin, an example of a classical ferritin enzyme, as a positive control. Unlike the Dps family of ferritin-like proteins, EncFtn showed no peroxidase activity when assayed with the substrate orho-phenylenediamine^25^. The ferroxidase activity of EncFtn_sH_ was measured by recording the progress curve of Fe^2+^ oxidation to Fe^3+^ at 315 nm after addition of 20 and 100 μM Fe^2+^ (2 and 10 times molar ratio Fe^2+^/FOC). In both experiments the rate of oxidation was faster than background oxidation of Fe^2+^ by molecular oxygen, and was highest for 100 μM Fe^2+^ (Figure 8A). This data showed that recombinant EncFtn_sH_ acts as an active ferroxidase enzyme. When compared to apoferritin, EncFtn_sH_ oxidized Fe^2+^ at a slower rate and the reaction did not run to completion over the 1800 seconds of the experiment. Addition of higher quantities of iron resulted in the formation of a yellow/red precipitate at the end of the reaction. We also performed these assays on purified recombinant encapsulin; which, when assayed alone, did not display ferroxidase activity above background Fe^2+^ oxidation (Figure 8B). In contrast, complexes of the full EncFtn encapsulin nanocompartment (i.e. the EncFtn-Enc protein complex) displayed ferroxidase activity comparable to apoferritin without the formation of precipitates (Figure 8B).

We attributed the precipitates observed in the EncFtn_sH_ ferroxidase assay to the production of insoluble Fe^3+^ complexes, which led us to propose that EncFtn does not directly store Fe^3+^ in a mineral form. This observation agrees with native MS results, which denotes a maximum iron loading of 10-15 iron ions per decameric EncFtn; and the structure, which does not possess the enclosed iron storage cavity characteristic of classical ferritins and Dps family proteins that can directly accrue mineralized Fe^3+^ within their nanocompartment structures.

To analyze the products of these reactions and determine whether the EncFtn and encapsulin were able to store iron in a mineral form, we performed TEM on the reaction mixtures from the ferroxidase assay. The EncFtn_sH_ reaction mixture showed the formation of large, irregular electron-dense precipitates (Figure 8-figure supplement 1A). A similar distribution of particles was observed after addition of Fe^2+^ to the encapsulin protein (Figure 8-figure supplement 1B). In contrast, addition of Fe^2+^ to the EncFtn-Enc nanocompartment resulted in small, highly regular, electron dense particles of approximately 5 nm in diameter (Figure 8-figure supplement 1C); we interpret these observations as controlled mineralization of iron within the nanocompartment. Addition of Fe^2+^ to apoferritin resulted in a mixture of large particles and small (~2 nm) particles consistent with partial mineralization by the ferritin and some background oxidation of the iron (Figure 8-figure supplement 1D). Negative stain TEM of these samples revealed that upon addition of iron, the EncFtn_sH_ protein showed significant aggregation (Figure 8-figure supplement 1F); while the encapsulin, EncFtn-Enc system, and apoferritin are present as distinct nanocompartments without significant protein aggregation (Figure8-figure supplement 1G-I).

### Iron storage in encapsulin nanocompartments

The results of the ferroxidase assay and micrographs of the reaction products suggest that the oxidation and mineralization function of the classical ferritins are split between the EncFtn and encapsulin proteins, with the EncFtn acting as a ferroxidase and the encapsulin shell providing an environment and template for iron mineralization and storage. To investigate this further, we added Fe^2+^ at various concentrations to samples of apo-ferritin, EncFtn, isolated encapsulin, and the EncFtn-Enc protein complex, and subjected these samples to a ferrozine assay to quantify the amount of iron associated with the proteins after three hours of incubation. The maximum iron loading capacity of these systems was calculated as the quantity of iron per biological assembly (Figure 8C). In this assay, the EncFtn_sH_ decamer binds a maximum of around 48 iron ions before excess iron induces protein precipitation. The encapsulin shell protein can sequester about 2200 iron ions before significant protein loss occurs, and the reconstituted EncFtn-Enc nanocompartment sequestered about 4150 iron ions. This latter result is significantly more than the apoferritin used in our assay, which sequesters approximately 570 iron ions over the same time period (Figure 8C, Table 5).

Consideration of the functional oligomeric states of these proteins, where EncFtn is a decamer and encapsulin forms an icosahedral cage, and estimation of the iron loading capacity of these complexes gives insight into the role of the two proteins in iron storage and mineralization. EncFtn decamers bind up to 48 iron ions (Fig 8C), which is significantly higher than the stoichiometry of fifteen metal ions visible in the FOC and E31/34-site of the crystal structure of the EncFtn_sH_ decamer and our MS analysis. The discrepancy between these solution measurements and our MS analysis may indicate that there are additional metal-binding sites on the interior channel and exterior faces of the protein; this is consistent with our identification of a number of weak metal-binding sites at the surface of the protein in the crystal structure (Figure 5D). These observations are consistent with hydrated Fe^2+^ ions being channeled to the active site from the E31/34-site and the subsequent exit of Fe^3+^ products on the outer surface, as is seen in other ferritin family proteins^25,26^. While the isolated encapsulin shell does not display any ferroxidase activity, it binds around 2000 iron ions in our assay (Table 5). This implies that the shell can bind a significant amount of iron on its outer and inner surfaces. While the maximum reported loading capacity of classical ferritins is approximately 4000 iron ions^21^, in our assay system we were only able to load apoferritin with around 570 iron ions. However, the recombinant EncFtn-Enc nanocompartment was able to bind over 4100 iron ions in the same time period, over seven times the amount seen for the apoferritin. We note that in our experimental system we do not reach the experimental maximum for apoferritin and therefore the total iron-loading capacity of our system may be significantly higher than in this experimental system.

Taken together, our data show that EncFtn can catalytically reduce Fe^2+^ to Fe^3+^; however, iron binding in EncFtn is limited to the FOC and several weaker surface metal binding sites. In contrast, the encapsulin protein displays no catalytic activity, but has the ability to bind a considerable amount of iron. Finally, the EncFtn-Enc nanocompartment complex retains the catalytic activity of EncFtn, and sequesters iron within the encapsulin shell at a higher level than the isolated components of the system, and at a significantly higher level than the classical ferritins^15^. Furthermore, our recombinant nanocompartments may not have the physiological subunit stoichiometry, and the iron-loading capacity of native nanocompartments is potentially much higher than the level we have observed.

### Mutagenesis of the EncFtn_sH_ Ferroxidase center

To investigate the structural and biochemical role played by the metal binding residues in the diiron FOC of EncFtn_sH_ we produced alanine mutations in each of these residues: Glu32, Glu62, and His65. These EncFtn_sH_ mutants were produced in *E. coli* cells grown in M9 minimal medium, both in the absence and presence of additional iron. The E32A and E62A mutants eluted from SEC at a volume consistent with the decameric form of EncFtn_sH_, with a small proportion of monomer; the H65A mutant eluted at a volume consistent with the monomeric form of EncFtn_sH_ (Figure 9). For all of the mutants studied, no change in oligomerization state was apparent upon addition of Fe^2+^ *in vitro.*

In addition to SEC studies, native mass spectrometry of the apo-EncFtn_sH_ mutants was performed and compared with the wild-type apo-EncFtn_sH_ protein (Figure 10). As described above, the apo-EncFtn_sH_ has a charge state distribution consistent with an unstructured monomer, and decamer formation is only initiated upon addition of ferrous iron. Both the E32A mutant and E62A mutant displayed charge state distributions consistent with decamers, even in the absence of Fe^2+^. This gas-phase observation is consistent with SEC measurements, which indicate both of these variants were also decamers in solution. Thus it seems that these mutations allow the decamer to form in the absence of iron in the FOC. In contrast to the glutamic acid mutants, MS analysis of the H65A mutant is similar to wild-type apo-EncFtn_sH_ and is present as a monomer; interestingly a minor population of dimeric H65A was also observed.

We propose that the observed differences in the oligomerization state of the E32A and E62A mutants compared to wild-type are due to the changes in the electrostatic environment within the FOC. At neutral pH the glutamic acid residues are negatively charged, while the histidine residues are predominantly in their uncharged state. In the wild-type EncFtn_sH_ this leads to electrostatic repulsion between subunits in the absence of iron. Coordination of Fe^2+^ in this site stabilizes the dimer and reconstitutes the active FOC. The geometric arrangement of E32 and E62 in the FOC explains their behavior in solution and the gas phase, where they both favor the formation of decamers due to the loss of a repulsive negative charge. The FOC in the H65A mutant is destabilized through the loss of this metal coordinating residue and potential positive charge carrier, thus favoring the monomer in solution and the gas phase.

To understand the impact of the mutants on the organization and metal binding of the FOC, we determined the X-ray crystal structures of each of the EncFtn_sH_ mutants (See Table 4 for data collection and refinement statistics). The crystal packing of all of the mutants in this study is essentially isomorphous to the EncFtn_sH_ structure. All of the mutants display the same decameric arrangement in the crystals as the EncFtn_sH_ structure, and the monomers superimpose with an average RMSD_Cα_ of less than 0.2 Å.

Close inspection of the region of the protein around the FOC in each of the mutants highlights their effect on metal binding (Figure 11 and Figure 11-figure supplement 1 – 4). In the E32A mutant the position of the side chains of the remaining iron coordinating residues in the FOC is essentially unchanged, but the absence of the axial-metal coordinating ligand provided by the E32 side chain abrogates metal binding in this site. The E31/34-site also lacks metal, with the side chain of E31 rotated by 180°at the C_β_ in the absence of metal (Figure 11-figure supplement 1). The E62A mutant has a similar effect on the FOC to the E32A mutant, however the entry site still has a calcium ion coordinated between residues E31 and E34 (Figure 11-figure supplement 2). The H65A mutant diverges significantly from the wild type in the position of the residues E32 and Y39 in the FOC. E32 appears in either the original orientation as the wild type and coordinates Ca^2+^ in this position, or it is flipped by 180° at the C_β_, moving away from the coordinated calcium ion in the FOC. Y39 moves closer to Ca^2+^ compared to the wild-type and coordinates the calcium ion (Figure 11-figure supplement 3). A single calcium ion is present in the entry site of this mutant; however, E31 of one chain is rotated away from the metal ion and is not involved in coordination.

Taken together the results of our data show that these changes to the FOC of EncFtn still permit the formation of the decameric form of the protein. While the proteins all appear decameric in crystals, their solution and gas-phase behavior differs considerably and the mutants no longer show metal-dependent oligomerization. These results highlight the importance of metal coordination in the FOC for the stability and assembly of the EncFtn protein.

To address the question of how mutagenesis of the iron coordinating residues affects the enzymatic activity of the EncFtn_sH_ protein we recorded progress curves for the oxidation of Fe^2+^ to Fe^3+^ by the different mutants as before. Mutagenesis of E32A and H65A reduces the activity of EncFtn_sH_ by about 40%-55%; the E62A mutant completely abrogates activity, presumably through the loss of the bridging coordination for the formation of the di-nuclear iron center of the FOC (Figure 12). Collectively, the effect of mutating these residues in the FOC confirms the importance of the iron coordinating residues for the ferroxidase activity of the EncFtn_sH_ protein.

## Discussion

Our study reports on a new class of ferritin-like proteins (EncFtn), which are associated with bacterial encapsulin nanocompartments (Enc). By studying the EncFtn from *R. rubrum* we demonstrate that iron binding results in assembly of EncFtn decamers, which display a unique annular architecture. Despite a radically different quaternary structure to the classical ferritins, the four-helical bundle scaffold and FOC of EncFtn_sH_ are strikingly similar to ferritin (Figure 6A). A sequence-based phylogenetic tree for proteins in the ferritin family was constructed; in addition to the classical ferritins, bacterioferritins and Dps proteins, our analysis included the encapsulin-associated ferritin-like proteins (Enc-Ftns) and a group related to these, but lacking the encapsulin sequence (Non-EncFtn). The analysis revealed that the Enc-Ftn and Non-EncFtn proteins form groups distinct from the other clearly delineated groups of ferritins, and represent outliers in the tree (Figure 13). While it is difficult to infer ancestral lineages in protein families, the similarity seen in the active site scaffold of these proteins highlights a shared evolutionary relationship between EncFtn proteins and other members of the ferritin superfamily that has been noted in previous studies^15,27^. From this analysis, we propose that the four-helical fold of the classical ferritins may have arisen through gene duplication of an ancestor of EncFtn. This gene duplication would result in the C-terminal region of one EncFtn monomer being linked to the N-terminus of another and thus stabilizing the four-helix bundle fold within a single polypeptide chain (Figure 6B). Linking the protein together in this way relaxes the requirement for the maintenance of a symmetrical FOC and thus provides a path to the diversity in active-site residues seen across the ferritin family (Figure 6A, residues E95, Q128 and E131 in PmFtn, Supplementary file 1)^15,27^.

### Relationship between ferritin structure and activity

The quaternary arrangement of classical ferritins into an octahedral nanocage and Dps into a dodecamer is absolutely required for their function as iron storage compartments^28^. The oxidation and mineralization of iron must be spatially separated from the host cytosol to prevent the formation of damaging hydroxyl radicals in the Fenton and Haber-Weiss reactions^29^. This is achieved in all ferritins by confining the oxidation of iron to the interior of the protein complex, thus achieving sequestration of the Fe^3+^ mineralization product. A structural alignment of the FOC of EncFtn with the classical ferritin PmFtn shows that the central ring of EncFtn corresponds to the external surface of ferritin, while the outer circumference of EncFtn is congruent with the inner mineralization surface of ferritin (Figure 6-figure supplement 1A). This overlay highlights the fact that the ferroxidase center of EncFtn faces in the opposite direction relative to the classical ferritins and is essentially inside out regarding iron storage space (Figure 6-figure supplement 1B, boxed region). Analysis of the point mutants made in the FOC highlights the importance of the iron-coordinating residues in the catalytic activity of EncFtn. Furthermore, the position of the calcium ion coordinated by E31 and E34 seen in the EncFtn_sH_ structure suggests an entry site to channel metal ions into the FOC; we propose that this site binds hydrated iron ions *in vivo* and acts as a selectivity filter and gate for the FOC^30^. The constellation of charged residues on the outer circumference of EncFtn (H57, E61 and E64) are consistent with residues lining the mineralization surface within the classical ferritin nanocage^18^, and given their proximity to the FOC these sites may be the exit portal and mineralization site^31^.

The absolute requirement for the spatial separation of oxidation and mineralization in ferritins suggests that the EncFtn family proteins are not capable of storing iron minerals due to the absence of an enclosed compartment in their structure (Figure 6-figure supplement 1B). Our biochemical characterization of EncFtn supports this hypothesis, indicating that while this protein is capable of oxidizing iron, it does not accrue mineralized iron in an analogous manner to classical ferritins. While EncFtn does not store iron itself, its association with the encapsulin nanocage suggests that mineralization occurs within the cavity of the encapsulin shell^3^. Our ferroxidase assay data on the recombinant EncFtn-Enc nanocompartments, which accrue over 4100 iron ions per complex and form regular nanoparticles, is consistent with the encapsulin protein acting as the store for iron oxidized by the EncFtn enzyme. TEM analysis of the reaction products shows the production of homogeneous iron nanoparticles only in the EncFtn-Enc nanocompartment (Figure 8-figure supplement 1).

Docking the decamer structure of EncFtn_sH_ into the pentamer of the *T. maritima* encapsulin Tmari_0786 (PDB ID: 3DKT)^1^ shows that the position of the C-terminal extensions of our EncFtn_sH_ structure are consistent with the localization sequences seen bound to the encapsulin protein (Figure 14A). Thus, it appears that the EncFtn decamer is the physiological state of this protein. This arrangement positions the central ring of EncFtn directly above the pore at the five-fold symmetry axis of the encapsulin shell and highlights a potential route for the entry of iron into the encapsulin and towards the active site of EncFtn. A comparison of the encapsulin nanocompartment and the ferritin nanocage highlights the size differential between the two complexes (Figure 14B) that allows the encapsulin to store significantly more iron. The presence of five FOCs per EncFtn_sH_ decamer and the fact that the icosahedral encapsulin nanocage can hold up to twelve of these complexes between each of the internal five-fold vertices means that they can achieve a high rate of iron mineralization across the entire nanocompartment. This arrangement of multiple reaction centers in a single protein assembly is reminiscent of classical ferritins, which has 24 FOCs distributed around the nanocage.

Our structural data, coupled with biochemical and ICP-MS analysis, suggest a model for the activity of the encapsulin iron-megastore (Figure 14C). The crystal structure of the *T. maritima* encapsulin shell protein has a negatively charged pore positioned to allow the passage of Fe^2+^ into the encapsulin and directs the metal towards the central, negatively charged hole of the EncFtn ring (Figure4-figure supplement 1). The metal binding sites on the interior of the ring (E31/34-sites) select for iron and direct it towards the five FOCs. We propose that the oxidation of Fe^2+^ to Fe^3+^ occurs within the FOC according to the model postulated by Ebrahimi *et al*^32^, in which the FOC acts as a substrate site through which iron passes and is released on to weakly coordinating sites at the outer circumference of the protein (H57, E61 and E64), where it is able to form ferrihydrite minerals which can be safely deposited within the lumen of the encapsulin nanocompartment (Figure 14).

Here we show for the first time the structure of a new class of ferritin-like protein and demonstrate that it has an absolute requirement for compartmentalization within an encapsulin nanocage to act as an iron store. Further work on the EncFtn-Enc nanocompartment will establish the structural basis for the movement of iron through the encapsulin shell, the mechanism of iron oxidation by the EncFtn FOC and its subsequent storage in the lumen of the encapsulin nanocompartment.

## Figure Legends

**Figure 1.**
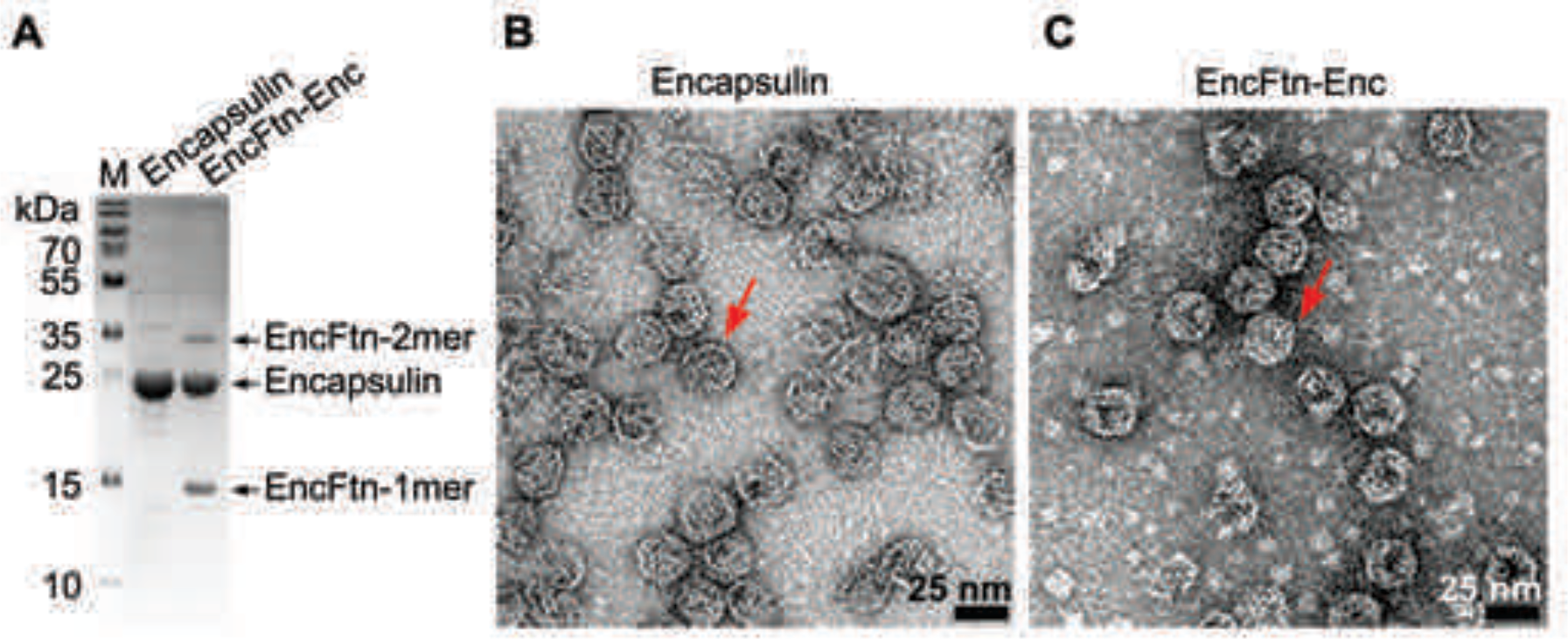
Purification of recombinant *Rhodospirillum rubrum* encapsulin nanocompartments. (**A**) Recombinantly expressed Encapsulin (Enc) and co-expressed EncFtn-Enc were purified by sucrose gradient ultracentrifugation from *E. coli* B834(DE3) grown in SeMet media. Samples were resolved by 18% acrylamide SDS-PAGE; the position of the proteins found in the complexes as resolved on the gel are shown with arrows. (**B/C**) Negative stain TEM image of recombinant encapsulin and EncFtn-Enc nanocompartments. Samples were imaged at 143,000 × magnification, with scale bar shown as 25 nm. Representative encapsulin and EncFtn-Enc complexes are indicated with red arrows.

**Figure 1-figure supplement 1.**
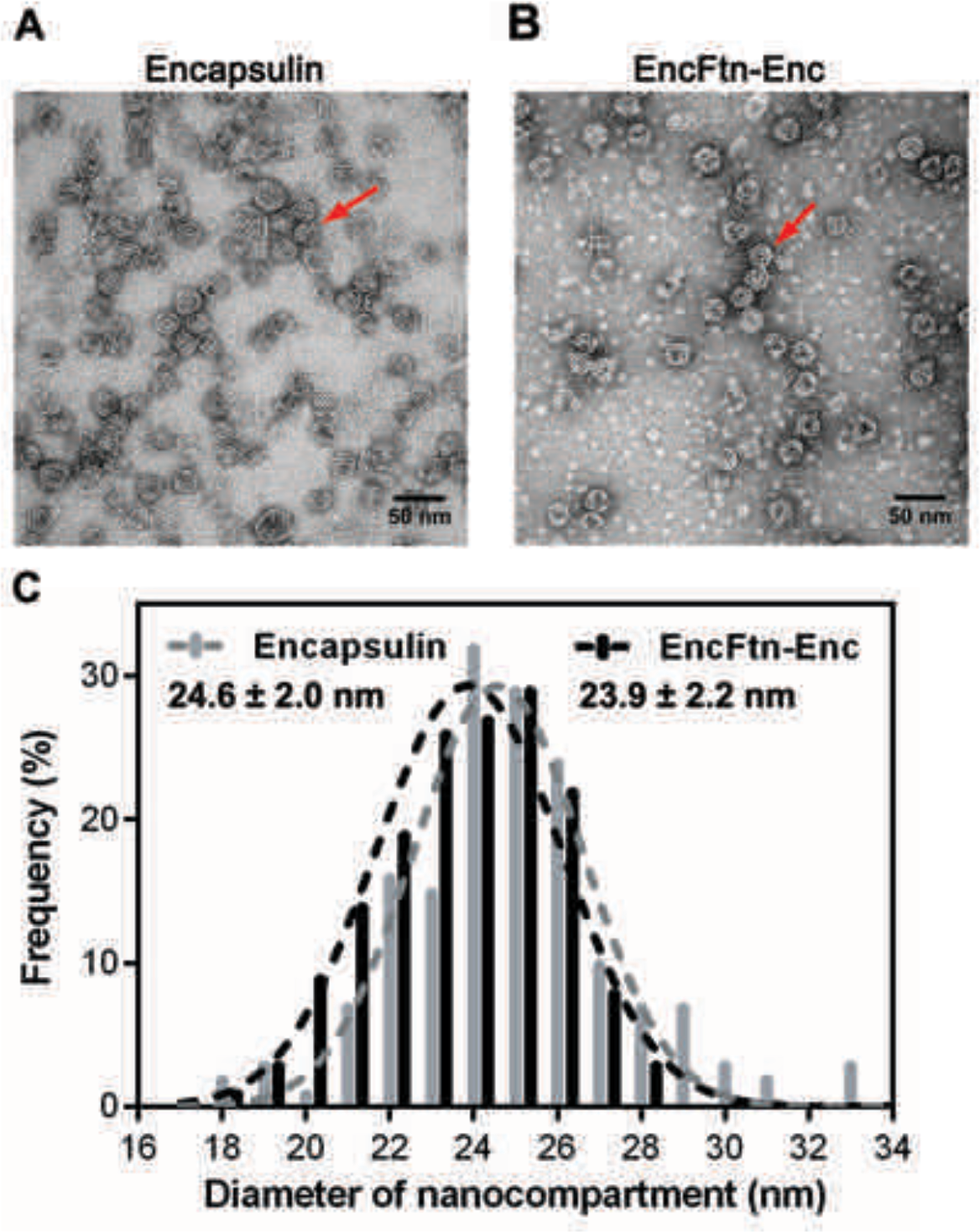
Full-frame transmission electron micrographs of *Rhodospirillum rubrum* nanocompartments. (**A/B**) Negative stain TEM image of recombinant *R. rubrum* encapsulin and EncFtn-Enc nanocompartments. All samples were imaged at 143,000 *X* magnification; the scale bar length corresponds to 50 nm. (**C**) Histogram showing the distribution of nanocompartment diameters. A model Gaussian nonlinear least square function was fitted to the data to obtain a mean diameter of 24.6 nm with SD of 2.0 nm for encapsulin (grey) and a mean value of 23.9 nm with SD of 2.2 nm for co-expressed EncFtn and encapsulin (EncFtn-Enc, black).

**Figure 2.**
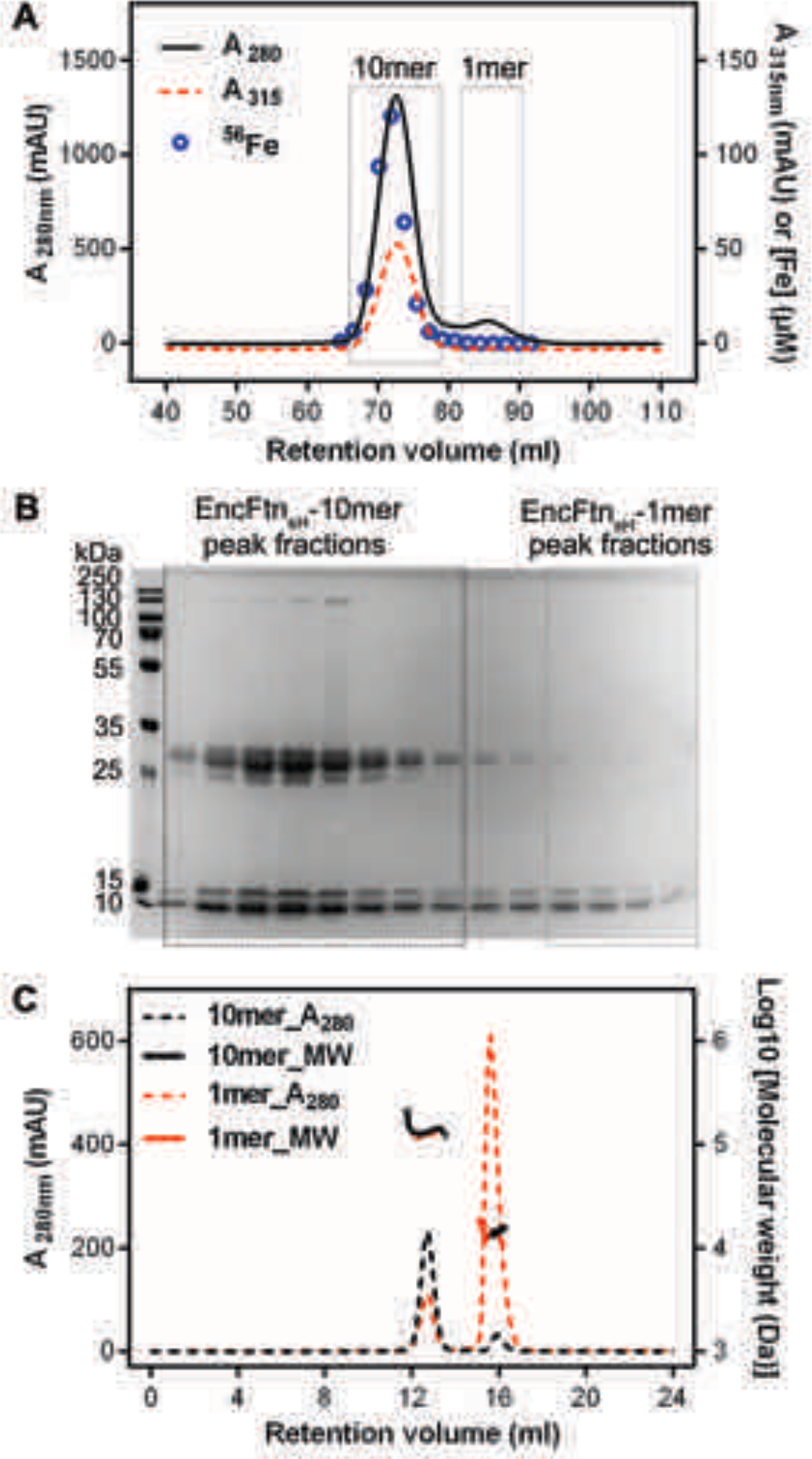
Purification of recombinant *Rhodospirillum rubrum* EncFtn_sH_. (**A**) Recombinant SeMet-labeled EncFtn_sH_ produced with 1 mM Fe(NH_4_)_2_(SO_4_)_2_ in the growth medium was purified by nickel affinity chromatography and size-exclusion chromatography using a S200 16/60 column (GE Healthcare). Chromatogram traces measured at 280 nm and 315 nm are shown with the results from ICP-MS analysis of the iron content of the fractions collected during the experiment. The peak around 73 ml corresponds to a molecular weight of around 130 kDa when compared to calibration standards; this is consistent with a decamer of EncFtn_sH_. The small peak at 85 ml corresponds to the 13 kDa monomer compared to the standards. Only the decamer peak contains significant amounts of iron as indicated by the ICP-MS analysis. (**B**) Peak fractions from the gel filtration run were resolved by 15% acrylamide SDS-PAGE and stained with Coomassie blue stain. The bands around 13 kDa and 26 kDa correspond to EncFtn_sH_, as identified by MALDI peptide mass fingerprinting. The band at 13 kDa is consistent with the monomer mass, while the band at 26 kDa is consistent with a dimer of EncFtn_sH_. The dimer species only appears in the decamer fractions. (C) SEC-MALLS analysis of EncFtn_sH_ from decamer fractions and monomer fractions allows assignment of an average mass of 132 kDa to decamer fractions and 13 kDa to monomer fractions, consistent with decamer and monomer species (Table 2).

**Figure 3.**
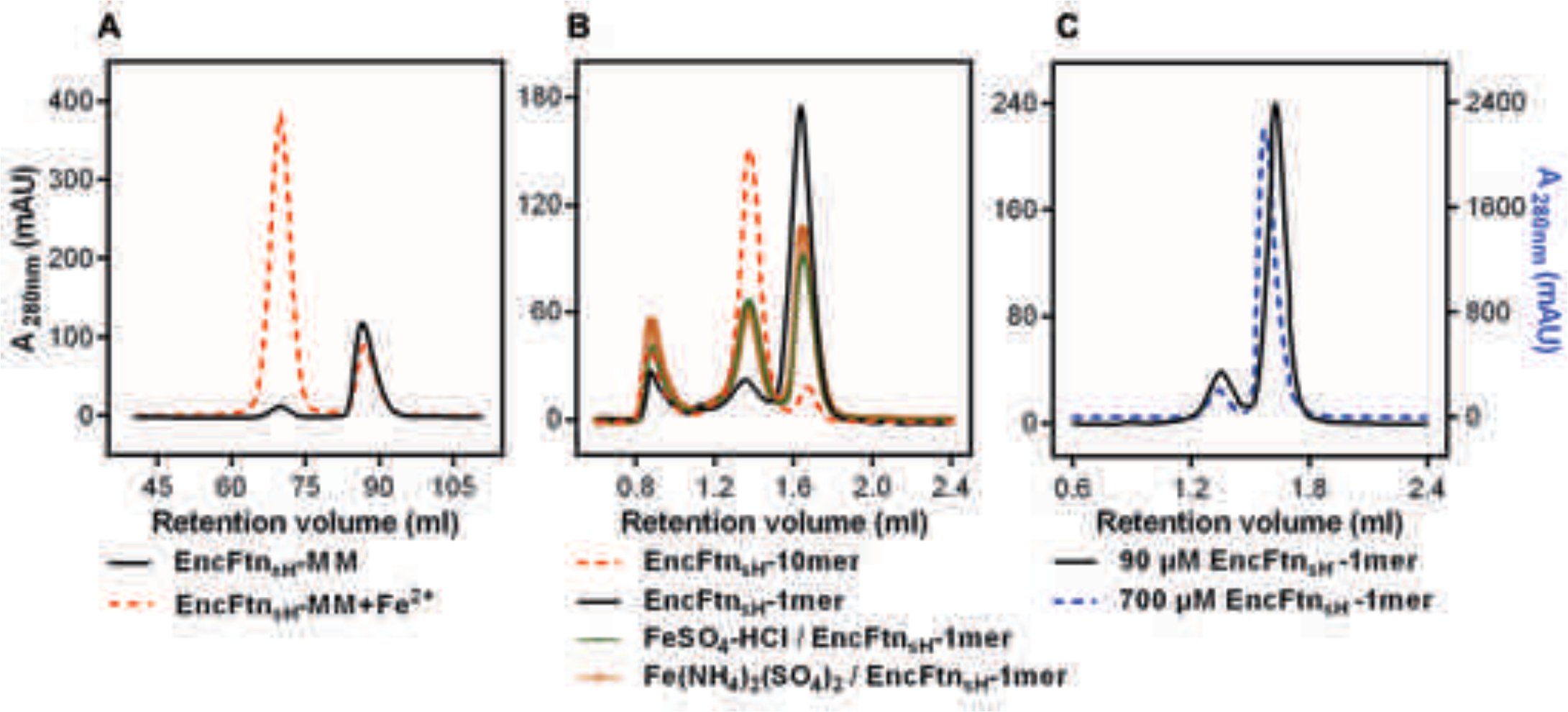
Effect of Fe^2+^ and protein concentration on the oligomeric state of EncFtn_sH_ in solution. (**A**) Recombinant EncFtn_sH_ was purified by preparative size-exclusion gel-filtration chromatography from *E. coli* BL21(DE3) grown in minimal media (MM) or in MM supplemented with 1 mM Fe(NH_4_)_2_(SO_4_)_2_ (MM+Fe^2+^). A higher proportion of decamer (peak between 65 and 75 ml) is seen in the sample purified from MM+Fe^2+^ compared to EncFtn_sH_-MM, indicating that Fe^2+^ facilitates the multimerization of EncFtn_sH_ *in vivo.* (**B**) EncFtn_sH_-monomer was incubated with one molar equivalent of Fe^2+^ salts for two hours prior to analytical gel-filtration using a Superdex 200 PC 3.2/30 column. Both Fe^2+^ salts tested induced the formation of decamer indicated by the peak between 1.2 and 1.6 ml. Monomeric and decameric samples of EncFtn_sH_ are shown as controls. Peaks around 0.8 ml were seen as protein aggregation. (**C**) Analytical gel filtration of EncFtn monomer at different concentrations to illustrate the effect of protein concentration on multimerization. The major peak shows a shift towards a dimer species at high concentration of protein, but the ratio of this peak (1.5 - 1.8 ml) to the decamer peak (1.2 - 1.5 ml) does not change when compared to the low concentration sample.

**Figure 3-figure supplement 1.**
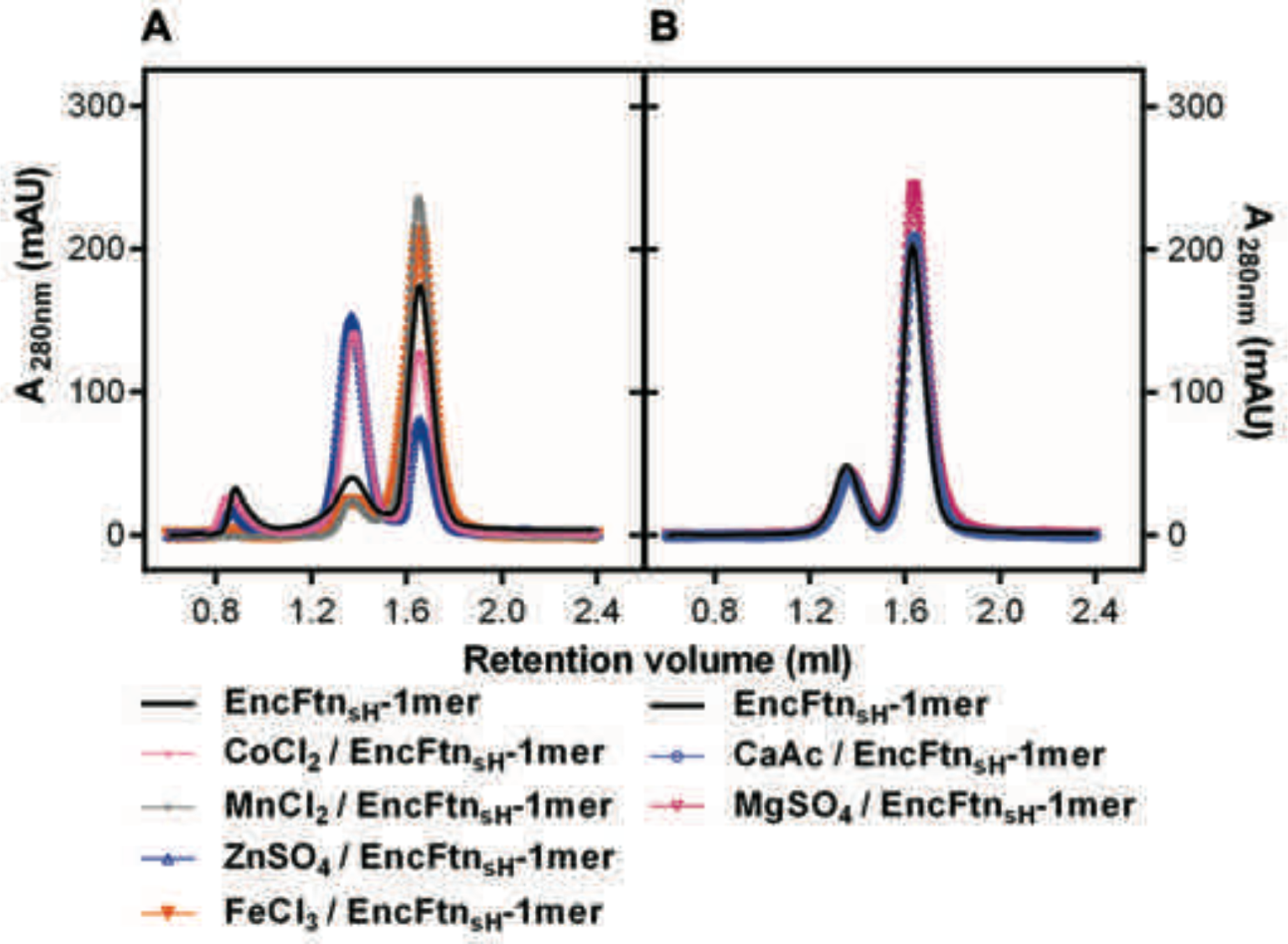
Effect of metal ions on the oligomeric state of EncFtn_sH_ in solution. (**A/B**) EncFtn_sH_-monomer was incubated with one mole equivalent of various metal salts for two hours prior to analytical gel-filtration using a Superdex 200 PC 3.2/30 column. Co^2+^ and Zn^2+^ induced the formation of the decameric form of EncFtn_sH_; while Mn^2+^, Mg^2+^ and Fe^3+^ did not significantly alter the oligomeric state of EncFtn_sH_.

**Figure 3-figure supplement 2.**
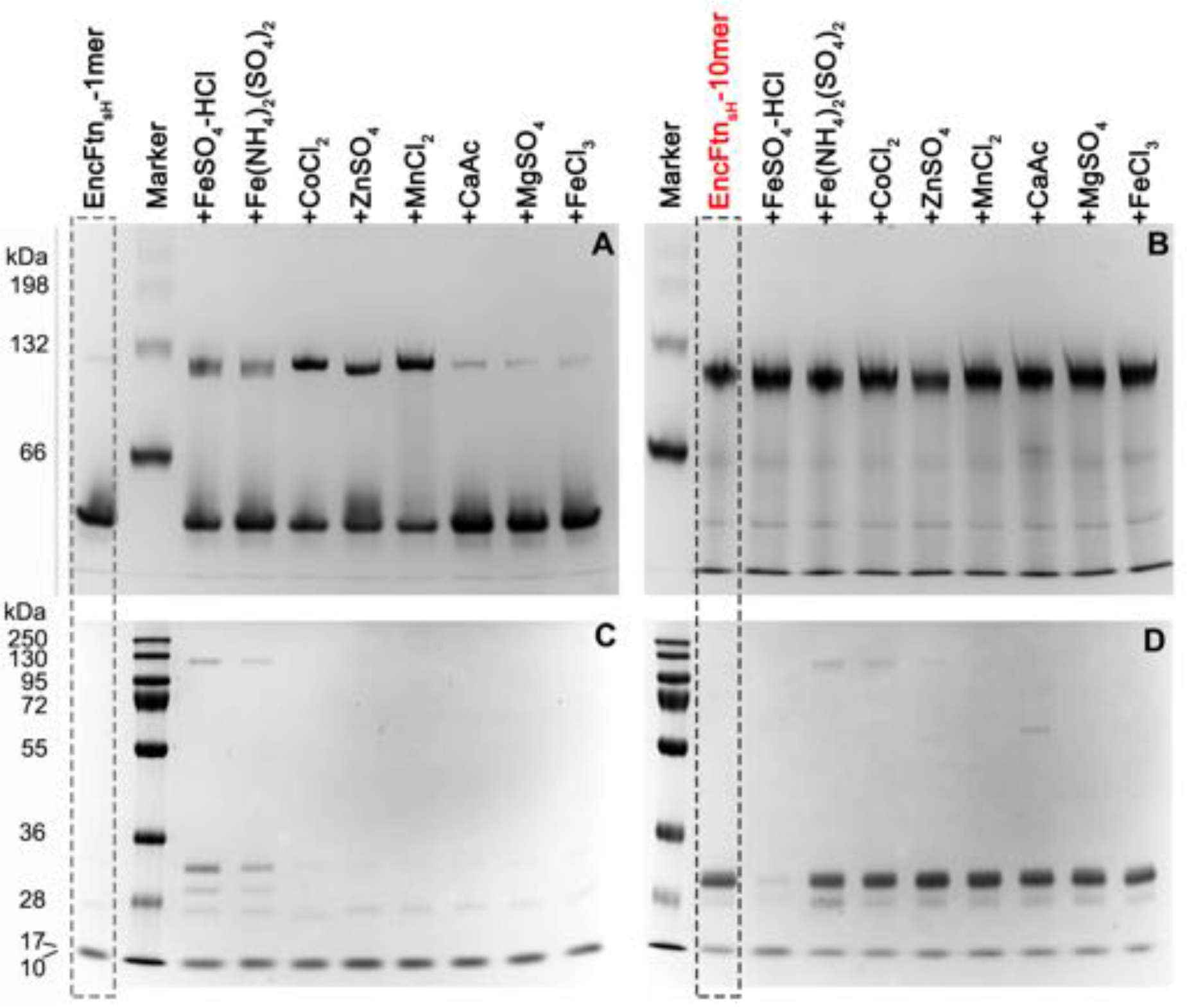
PAGE analysis of the effect of metal ions on the oligomeric state of EncFtn_sH_. 50 μM EncFtn_sH_ monomer or decamer samples were mixed with equal molar metal ions including Fe^2+^, Co^2+^, Zn^2+^, Mn^2+^, Ca^2+^, Mg^2+^ and Fe^3+^, which were analyzed by Native PAGE alongside SDS-PAGE. (**A**) 10% Native PAGE analysis of EncFtn_sH_ monomer fractions mixed with various metal solutions; (**B**) 10% Native PAGE analysis of EncFtn_sH_ decamer fractions mixed with various metal solutions; (**C**) 15% SDS-PAGE analysis on the mixtures of EncFtn_sH_ monomer fractions and metal solutions; (**D**) 15% SDS-PAGE analysis on the mixtures of EncFtn_sH_ decamer fractions and metal solutions.

**Figure 4.**
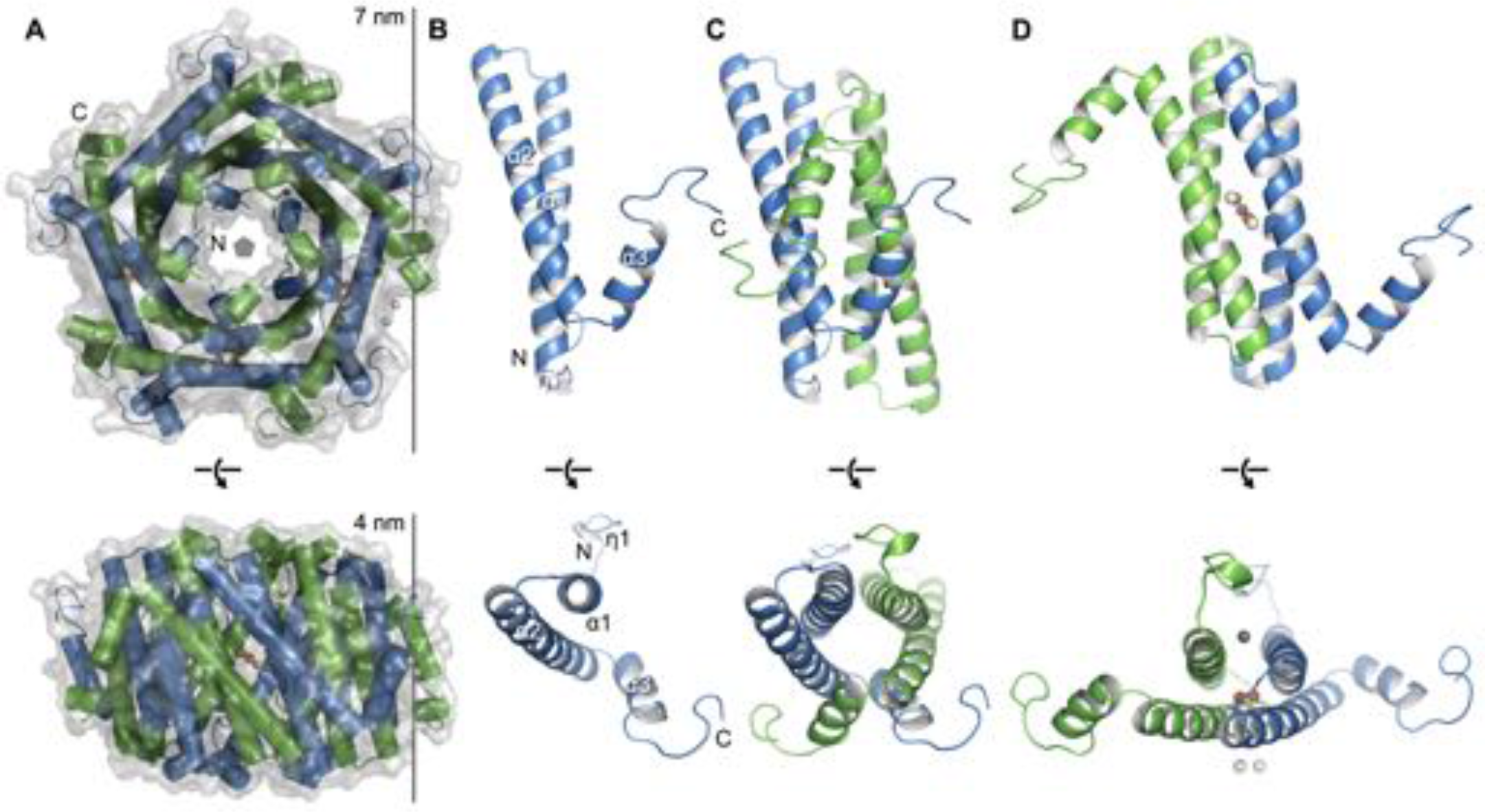
Crystal structure of EncFtn_sH_. (**A**) Overall architecture of EncFtn_sH_. Transparent solvent accessible surface view with α-helices shown as tubes and bound metal ions as spheres. Alternating subunits are colored blue and green for clarity. The doughnut-like decamer is 7 nm in diameter and 4.5 nm thick. (**B**) Monomer of EncFtn_sH_ shown as a secondary structure cartoon. (**C/D**) Dimer interfaces formed in the decameric ring of EncFtn_sH_. Subunits are shown as secondary structure cartoons and colored blue and green for clarity. Bound metal ions are shown as orange spheres for Fe^3+^ and grey and white spheres for Ca^2+^.

**Figure 4-figure supplement 1.**
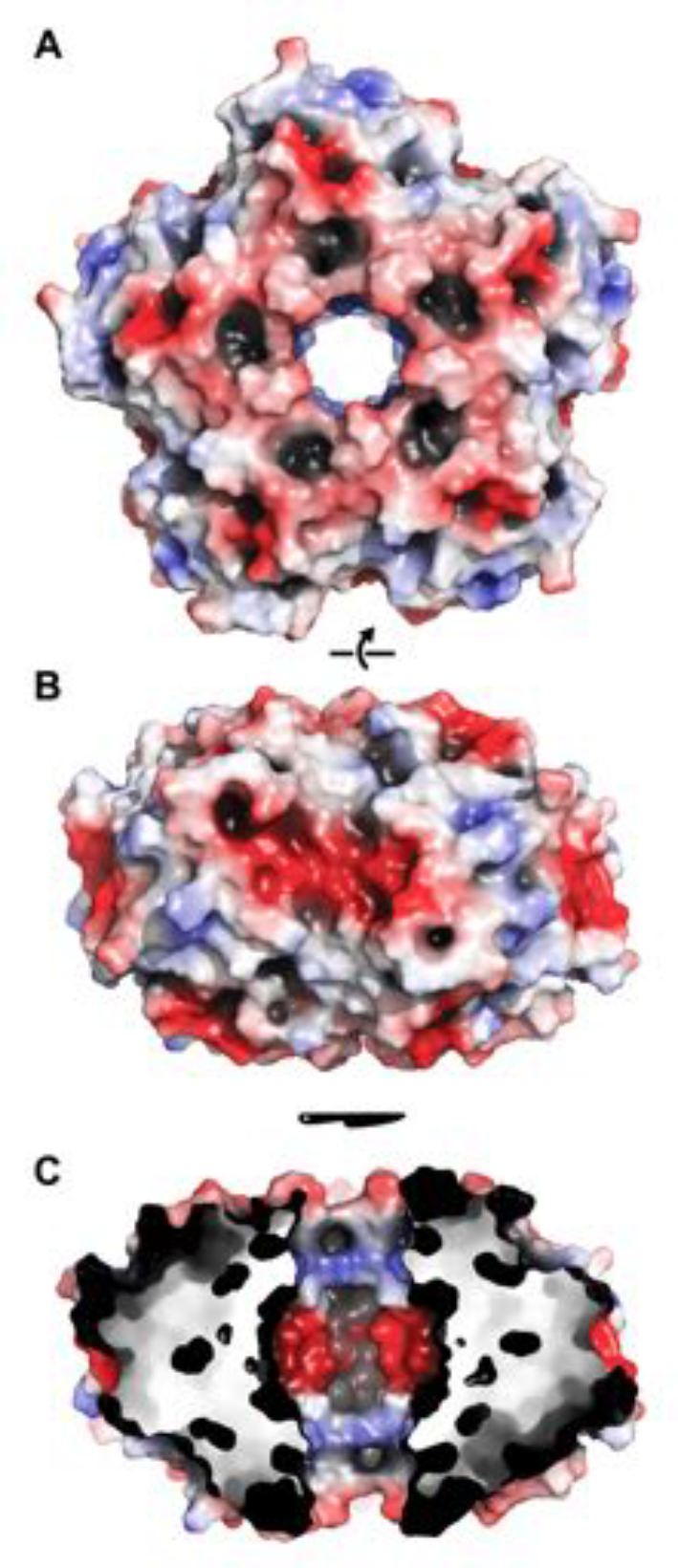
Electrostatic surface of EncFtn_sH_. The solvent accessible surface of EncFtn_sH_ is shown, colored by electrostatic potential as calculated using the APBS plugin in PyMol. Negatively charged regions are colored red and positive regions in blue, neutral regions in grey. (**A**) View of the surface of the EncFtn_sH_ decamer looking down the central axis. (**B**) Orthogonal view of (**A**). (**C**) Cutaway view of (**B**) showing the charge distribution within the central cavity.

**Figure 5.**
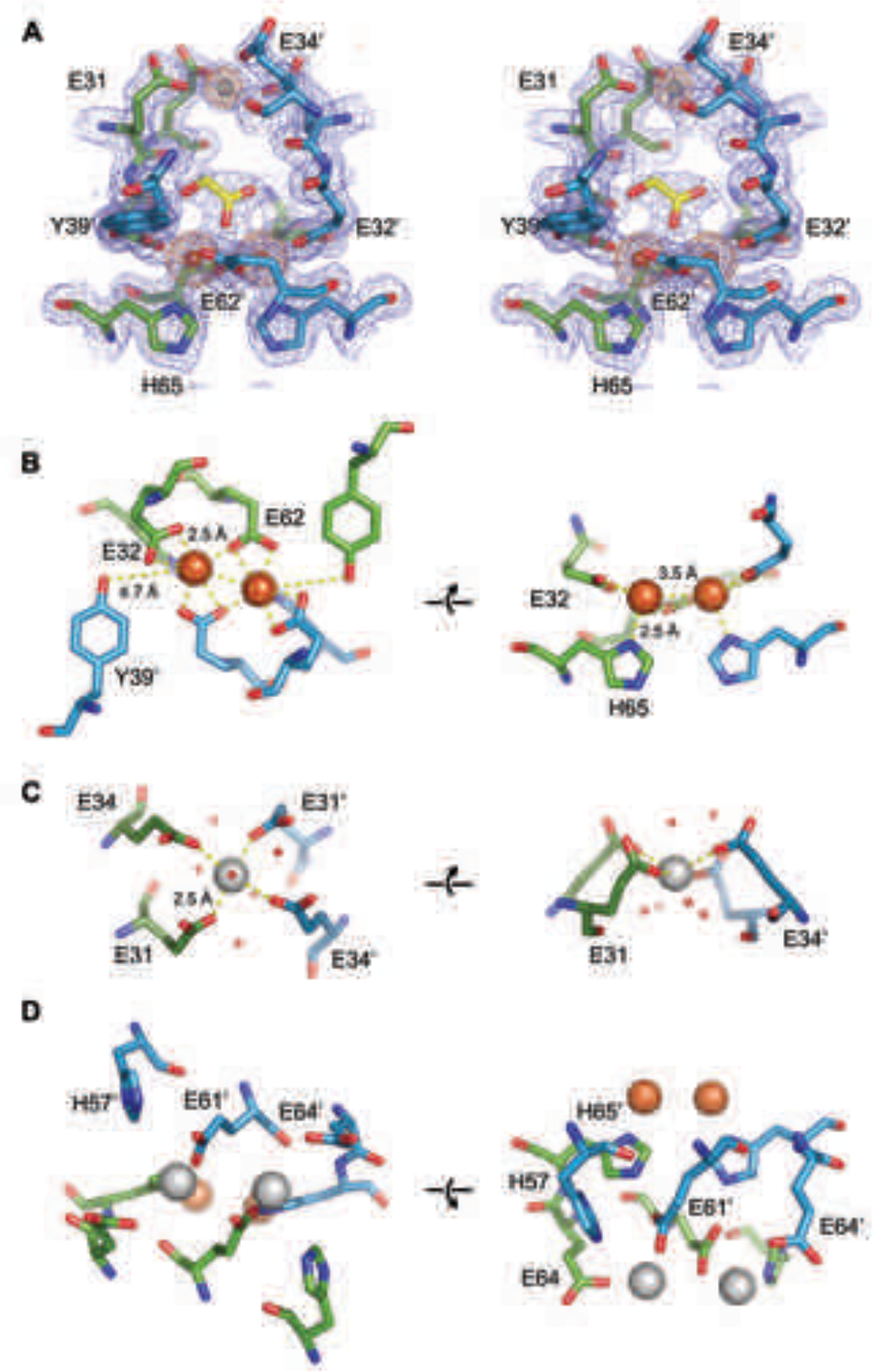
EncFtn_sH_ metal binding sites. (**A**) Wall-eyed stereo view of the metal-binding dimerization interface of EncFtn_sH_. Protein residues are shown as sticks with blue and green carbons for the different subunits, iron ions are shown as orange spheres and calcium as grey spheres, and the glycolic acid ligand is shown with yellow carbon atoms coordinated above the diiron center. The 2mFo-DFc electron density map is shown as a blue mesh contoured at 1.5 σ and the anomalous difference map is shown as an orange mesh and contoured at 10 σ. (**B**) Iron coordination within the FOC including residues E32, E62, H65 and Y39 from two chains. Protein and metal ions are shown as in (**A**). Coordination between the protein and iron ions is shown as yellow dashed lines with distances indicated. (**C**) Coordination of calcium within the dimer interface by four glutamic acid residues (E31 and E34 from two chains). The calcium ion is shown as a grey sphere and water molecules involved in the coordination of the calcium ion are shown as crosses. (**D**) Metal coordination site on the outer surface of EncFtn_sH_. The two calcium ions are coordinated by residues H57, E61 and E64 from the two chains of the FOC dimer, and are located at the outer surface of the complex, positioned 10 Å away from the FOC iron.

**Figure 5-figure supplement 1.**
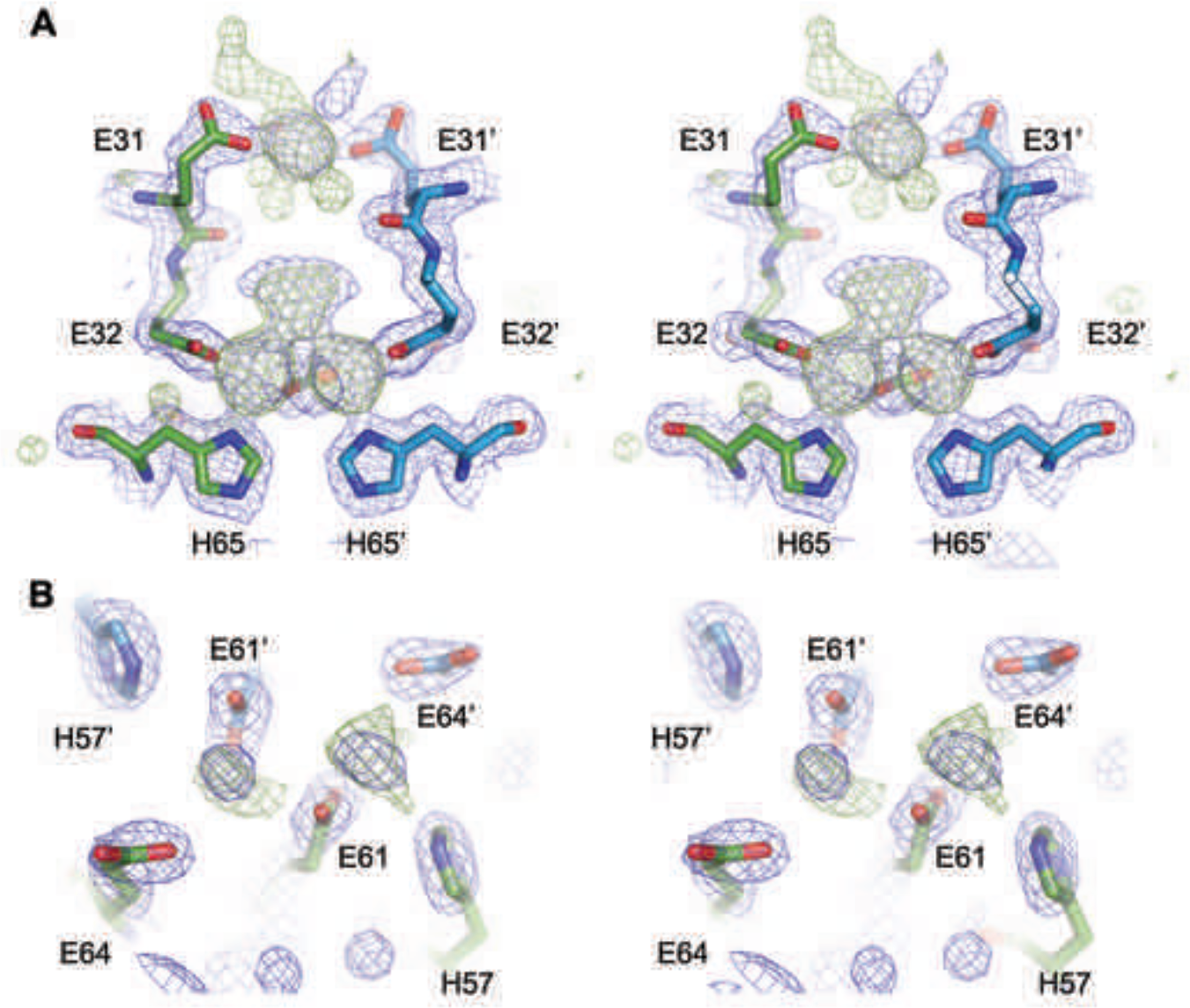
Putative ligand-binding site in EncFtn_sH_. (**A**) Wall-eyed stereo view of the dimer interface of EncFtn. Protein chains are shown as sticks, with 2mFo-DFc electron density shown in blue mesh and contoured at 1.5 σ and mFo-DFc shown in green mesh and contoured at 3 σ. (**B**) Wall-eyed stereo view of putative metal binding site at the external surface of EncFtn_sH_. Protein chains and electron density maps are shown as in (**A**).

**Figure 6.**
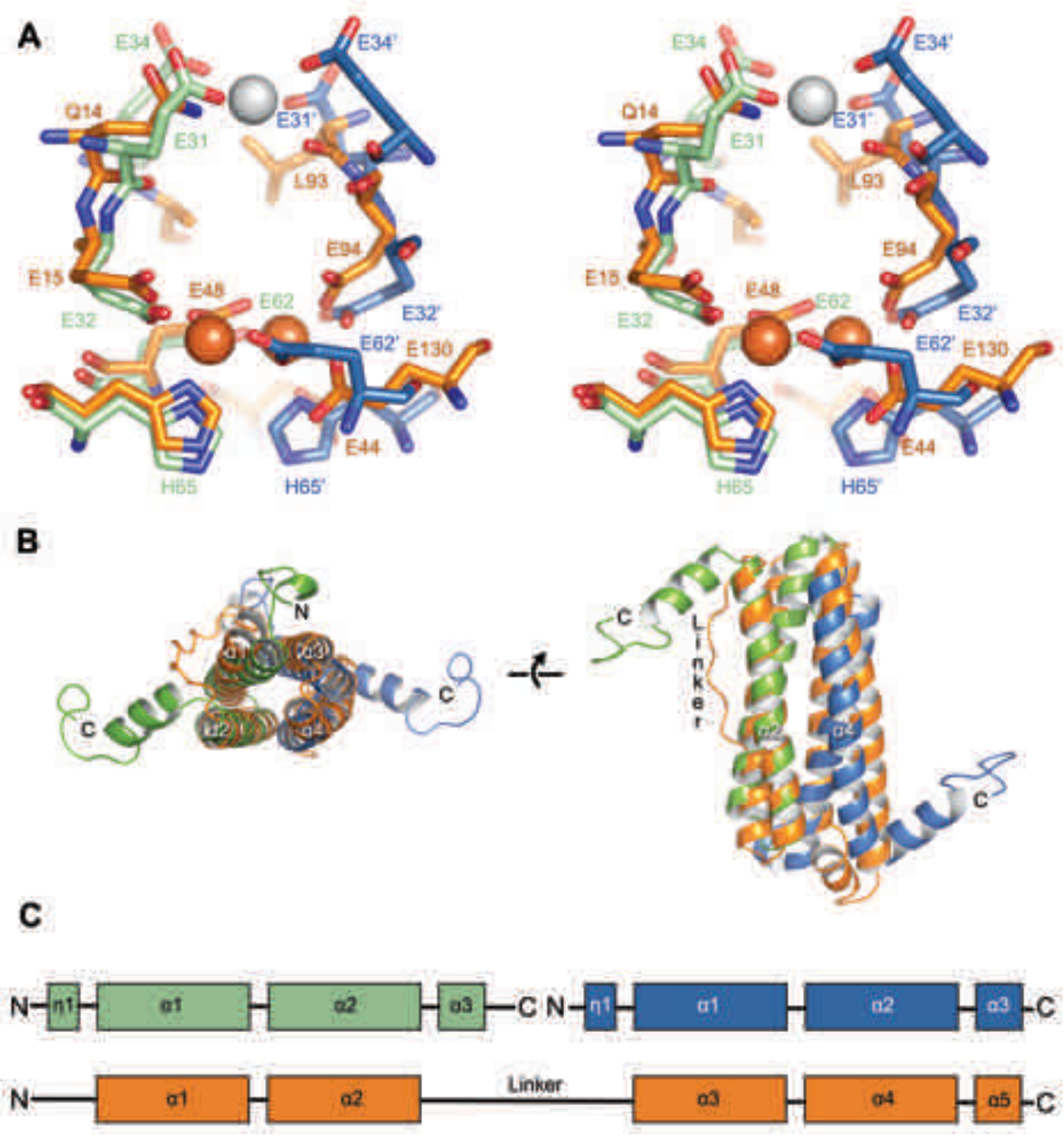
Comparison of the symmetric metal ion binding site of EncFtn_sH_ and the ferritin FOC. (**A**) Structural alignment of the FOC residues in a dimer of EncFtn_sH_ (green/blue) with a monomer of *Pseudo-nitzschia multiseries* ferritin (PmFtn) (PDBID: 4ITW) (orange)^33^. Iron ions are shown as orange spheres and a single calcium ion as a grey sphere. Residues within the FOC are conserved between EncFtn and ferritin PmFtn, with the exception of residues in the position equivalent to H65’ in the second subunit in the dimer (blue). The site in EncFtn with bound calcium is not present in other family members. (**B**) Secondary structure of aligned dimeric EncFtn_sH_ and monomeric ferritin highlighting the conserved four-helix bundle. EncFtn_sH_ monomers are shown in green and blue and aligned PmFtn monomer in orange as in A. (**C**) Cartoon of secondary structure elements in EncFtn dimer and ferritin. In the dimer of EncFtn that forms the FOC, the C-terminus of the first monomer (green) and N-terminus of the second monomer (blue) correspond to the position of the long linker between α2 and α3 in ferritin PmFtn.

**Figure 6-figure supplement 1.**
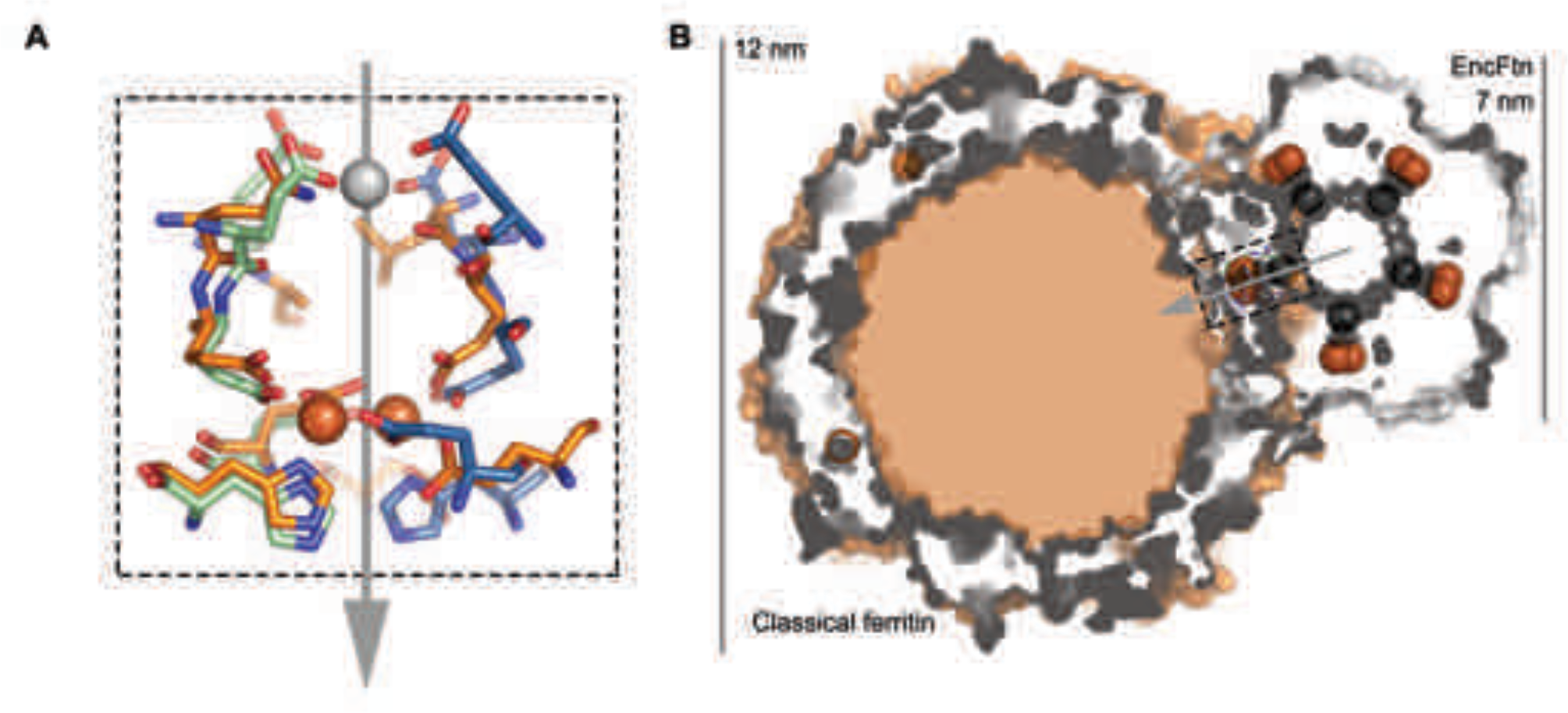
Comparison of quaternary structure of EncFtn_sH_ and ferritin. (**A**) Aligned FOC of EncFtn_sH_ and *Pseudo-nitzschia multiseries* ferritin (PmFtn)^33^. The metal binding site residues from two EncFtn_sH_ chains are shown in green and blue, while the PmFtn is shown in orange. Fe^2+^ in the FOC is shown as orange spheres and Ca^2+^ in EncFtn_sH_ is shown as a grey sphere. The 2-fold symmetry axis of the EncFtn FOC is shown with a grey arrow (**B**) Cross-section surface view of quaternary structure of EncFtn_sH_ and PmFtn as aligned in (**A**) (dashed black box). The central channel of EncFtn_sH_ is spatially equivalent to the outer surface of ferritin and its outer surface corresponds to the mineralization surface within ferritin.

**Figure 7.**
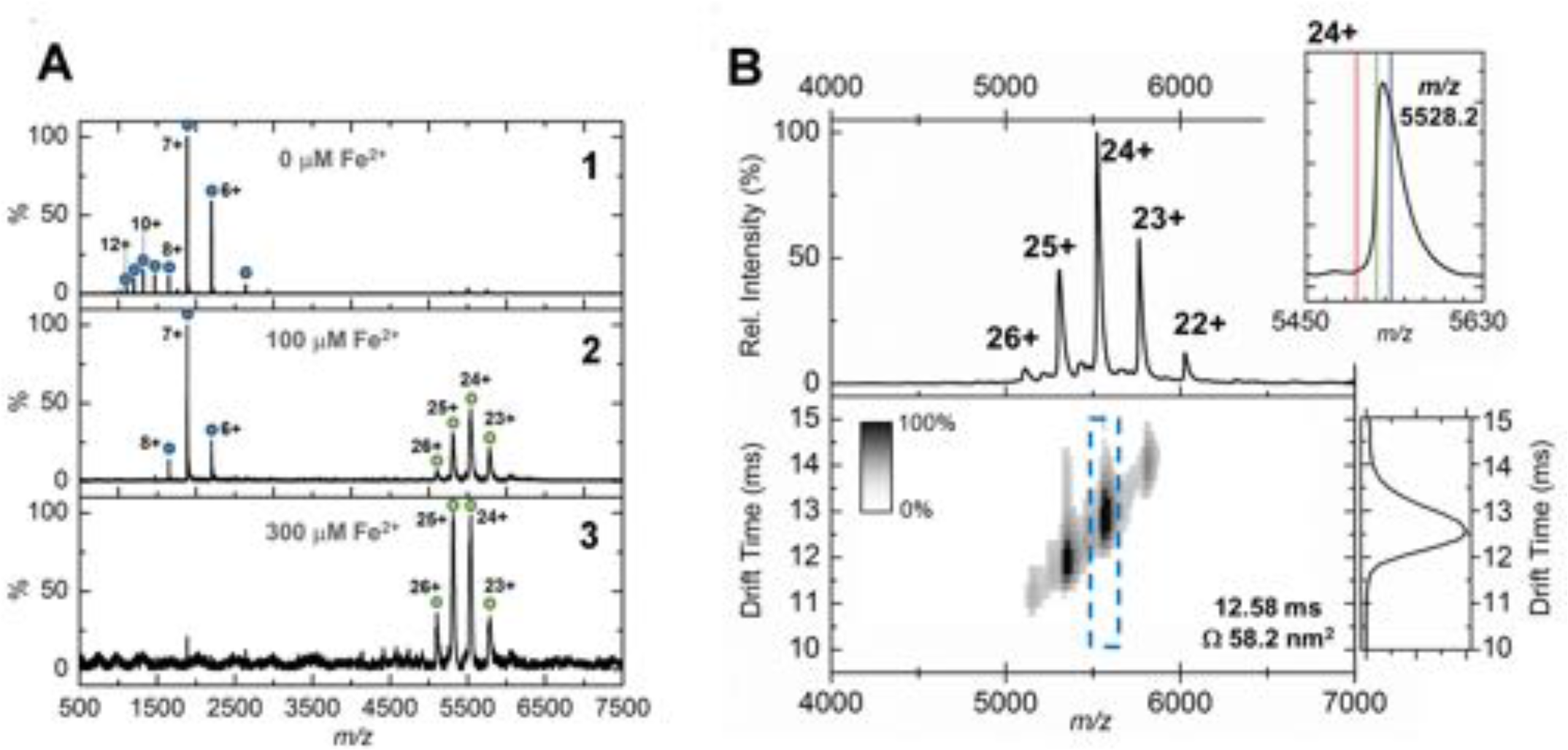
Native mass spectrometry and ion mobility analysis of iron loading in EncFtn_sH_. All spectra were acquired in 100 mM ammonium acetate, pH 8.0 with a protein concentration of 5 μM. (**A**) Native nanoelectrospray ionization (nESI) mass spectrometry of EncFtn_sH_ at varying iron concentrations. A1, nESI spectrum of iron-free EncFtn_sH_ displays a charge state distribution consistent with EncFtn_sH_ monomer (blue circles, 13,194 Da). Addition of 100 μM (A2) and 300 μM (A3) Fe^2+^ results in the appearance of a second higher molecular weight charge state distribution consistent with a decameric assembly of EncFtn_sH_ (green circles, 132.6 kDa). (**B**) Ion mobility (IM)- MS of the iron-bound holo-EncFtn_sH_ decamer. *Top*, Peaks corresponding to the +22 to +26 charge states of a homo-decameric assembly of EncFtn_sH_ are observed (132.6 kDa). *Top Insert*, Analysis of the 24+ charge state of the assembly at *m*/*z* 5528.2 Th. The theoretical average *m*/*z* of the 24+ charge state with no addition metals bound is marked by a red line (5498.7 Th); the observed *m*/*z* of the 24+ charge state indicates that the EncFtn_sH_ assembly binds between 10 (green line, 5521.1 Th) and 15 Fe ions (blue line, 5532.4 Th) per decamer. *Bottom*, the arrival time distributions (ion mobility data) of all ions in the EncFtn_sH_ charge state distribution displayed as a greyscale heat map (linear intensity scale). *Bottom right*, the arrival time distribution of the 24+ charge state (dashed blue box) has been extracted and plotted. The drift time for this ion is shown (ms), along with the calibrated collision cross section (CCS), Ω (nm^2^).

**Figure 7-figure supplement 1.**
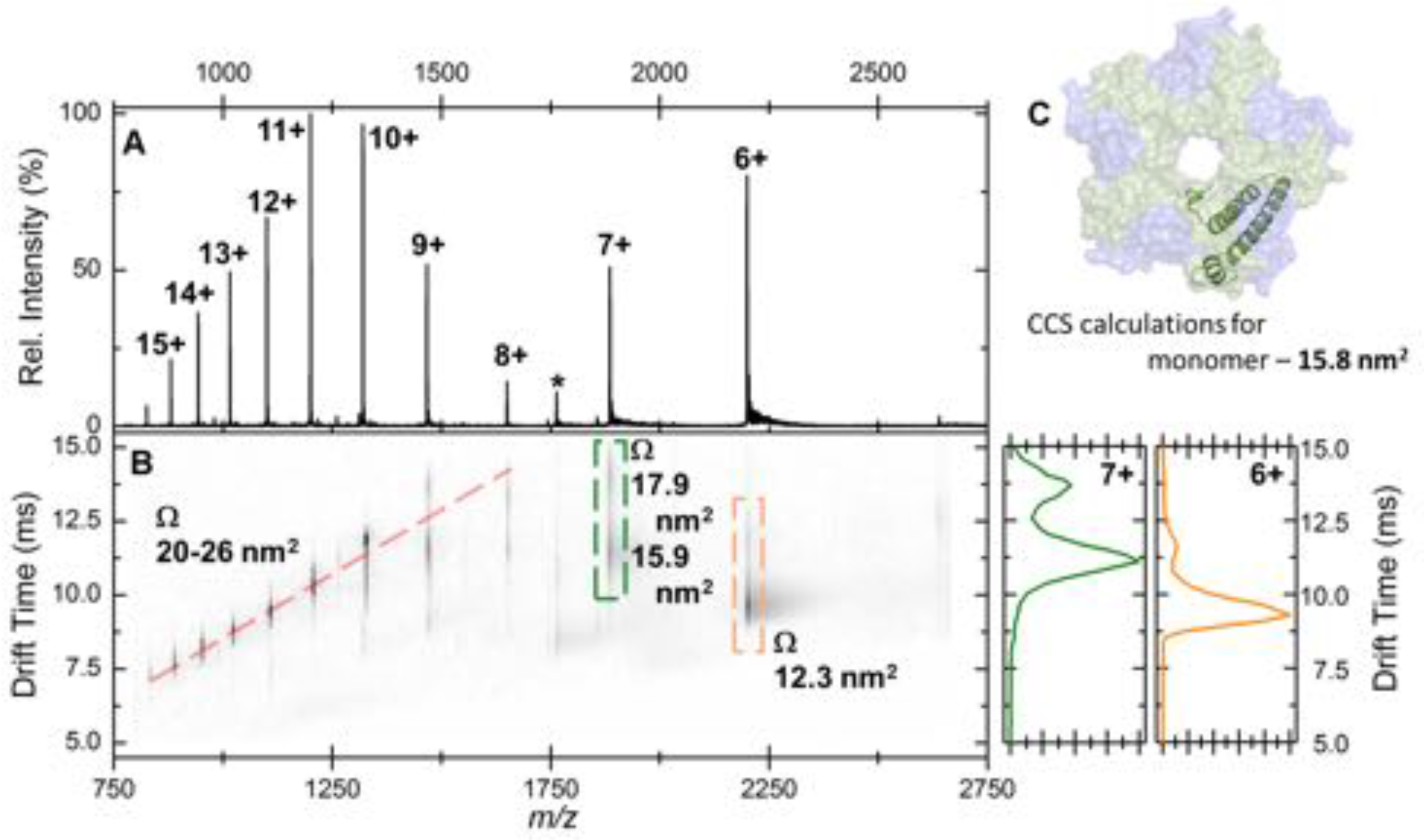
Native IM-MS analysis of the apo-EncFtn_sH_ monomer. (**A**) Mass spectrum of apo-EncFtn_sH_ acquired from 100 mM ammonium acetate pH 8.0 under native MS conditions. The charge state distribution observed is bimodal, with peaks corresponding to the 6+ to 15+ charge states of apo-monomer EncFtn_sH_ (neutral average mass 13,194.3 Da). (**B**), the arrival time distributions (ion mobility data) of all ions in the apo-EncFtn_sH_ charge state distribution displayed as a greyscale heat map (linear intensity scale). (**B**) ***right***, the arrival time distribution of the 6+ (orange) and 7+ (green) charge state (dashed colored-box) has been extracted and plotted; The arrival time distributions for these ion is shown (ms), along with the calibrated collision cross section, Ω (nm^2^). (**C**) The collision cross section of a single monomer unit from the crystal structure of the Fe-loaded EncFtn_sH_ decamer was calculated to be 15.8 nm^2^ using IMPACT v. 0.9.1. The +8 to +15 protein charge states have observed CCS between 20-26 nm^2^, which is significantly higher than the calculated CCS for an EncFtn_sH_ monomer taken from the decameric assembly crystal structure (15.8 nm). The mobility of the +7 charge state displays broad drift-time distribution with maxima consistent with CCS of 15.9 and 17.9 nm^2^. Finally, the +6 charge state of EncFtn_sH_ has mobility consistent with a CCS of 12.3 nm^2^, indicating a more compact / collapsed structure. It is clear from this data that apo-EncFtn_sH_ exists in several gas phase conformations. The range of charge states occupied by the protein (6+ to 15+) and the range of CCS in which the protein is observed (12.3 nm^2^ – 26 nm^2^) are both large. In addition, many of the charge states observed have higher charge than the theoretical maximal charge on spherical globular protein, as determined by the De La Mora relationship (Z_R_ = 0.0778√*m*; for the EncFtn_sH_ monomer Z_R_ = 8.9)^34^. As described by Beveridge et al., all these factors are indicative of a disordered protein^35^.

**Figure 7-figure supplement 2.**
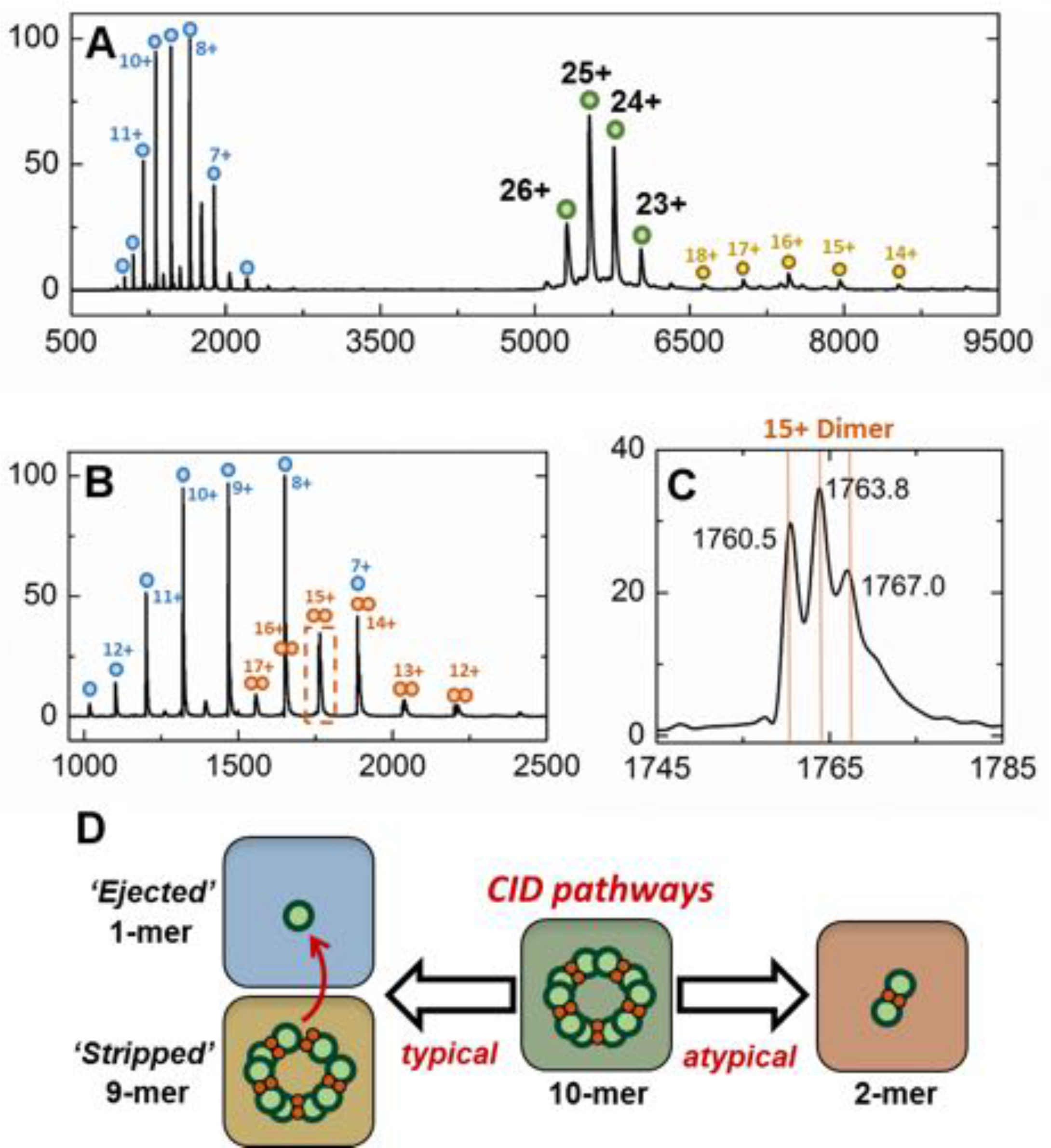
Gas-phase disassembly of the holo-EncFtn_sH_ decameric assembly. The entire charge state distribution of the Fe-loaded holo-EncFtn assembly (green circles) was subject to collisional-induced dissociation (CID) by increasing the source cone voltage to 200 V and the trap voltage to 50 V. The resulting CID mass spectrum (**A**) revealed that dissociation of the holo-EncFtn decamer primarily occurred via ejection of a highly charged monomer (blue circles), leaving the 'stripped’ complex (a 9mer; 118.7 kDa; yellow circles). The mass of the ejected-monomer is consistent with apo-EncFtn (13.2 kDa), suggesting unfolding of the monomer (and loss of Fe) occurs during ejection from the complex. This observation of asymmetric charge partitioning of the sub-complexes with respect to the mass of the complex is consistent with the “typical” pathway of dissociation of protein assemblies by CID, as described by Hall *et al*^36^. In addition, a third, lower abundance, charge state distribution is observed which overlaps the EncFtn ejected monomer charge state distribution; this region of the spectrum is highlighted in (**B**). This distribution is consistent with an ejected EncFtn dimer (orange circles). Interestingly, closer analysis of the individual charge state of this dimeric CID product shows that this sub-complex exists in three forms - displaying mass consistent with an EncFtn dimer binding 0, 1, and 2 Fe ions. This is highlighted in (**C**), where the 15+ charge state of the EncFtn dimer is shown; 3 peaks are observed with *m/z* 1760.5, 1763.8, and 1767.0 Th - the lowest peak corresponds to neutral masses of 26392.5 Da [predicted Enc-Ftn dimer, (C_572_H_884_N_172_O_185_S_2_)_2_ - 26388.6 Da]. The two further peaks have a delta-mass of ~+50 Da, consistent with Fe binding. We interpret these observations as partial ‘atypical’ CID fragmentation of the decameric complex - i.e. fragmentation of the initial complex with retention of subunit and ligand interactions. A schematic summary of these results is displayed in (**D**). We postulate the high stability of this iron-bound dimer sub-complex is due to the metal coordination at the dimer interface, increasing the strength of the dimer interface. Taken together, these observations support our findings that the topology of the decameric EncFtn_sH_ assembly is arranged as a pentamer of dimers, with two Fe ions at each dimer interface.

**Figure 8.**
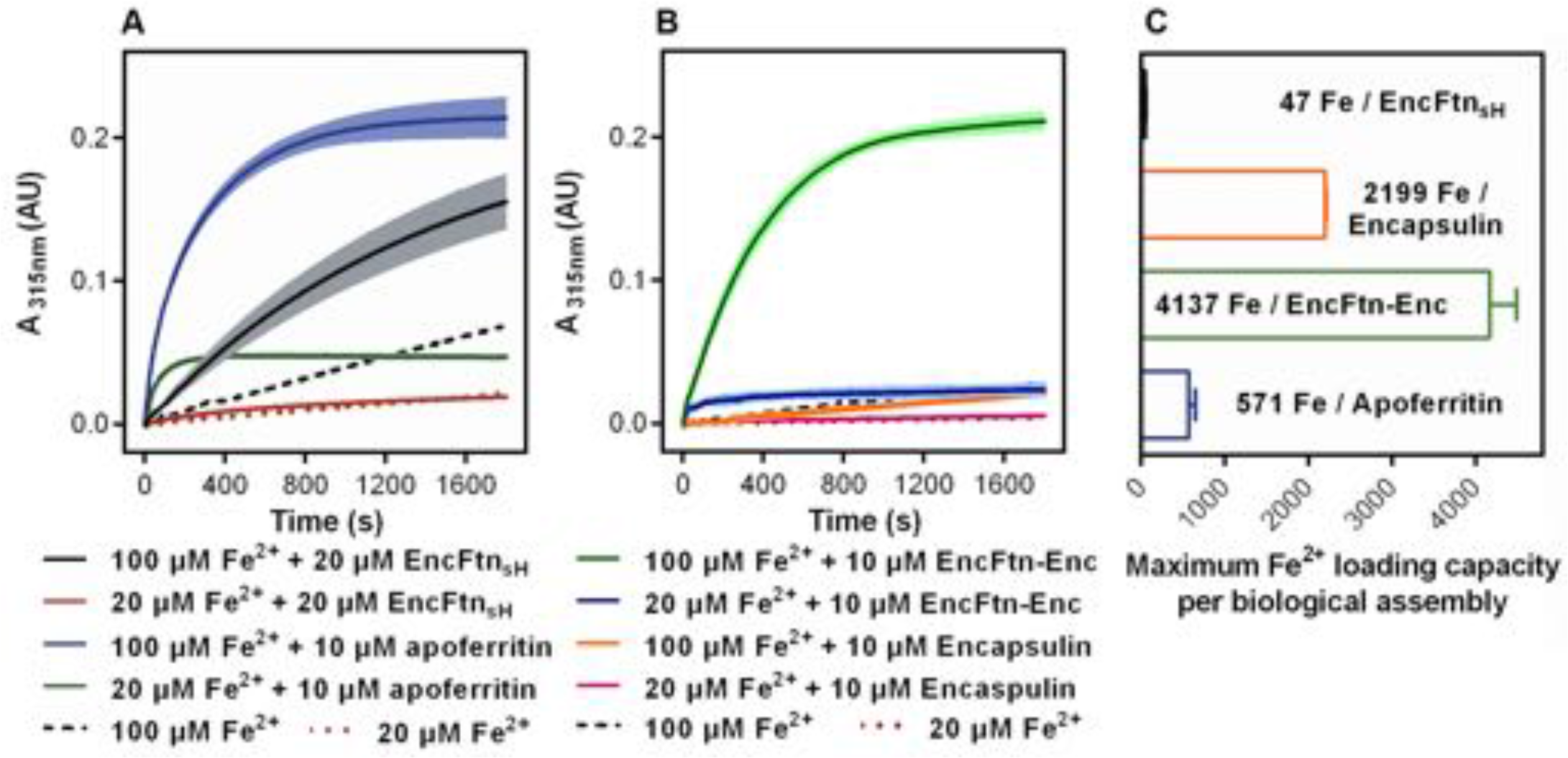
Spectroscopic evidence for the ferroxidase activity and comparison of iron loading capacity of apoferritin, EncFtn_sH_, encapsulin, and EncFtn-Enc. (**A**) Apoferritin (10 μM monomer concentration) and EncFtn_sH_ (20 μM monomer concentration, 10 μM FOC concentration) were incubated with 20 and 100 μM iron (2 and 10 times molar equivalent Fe^2+^ per FOC) and progress curves of the oxidation of Fe^2+^ to Fe^3+^ at 315 nm were recorded in a spectrophotometer. The background oxidation of iron at 20 and 100 μM in enzyme-free controls are shown for reference. (**B**) Encapsulin and EncFtn-Enc complexes at 10 μM asymmetric unit concentration were incubated with Fe^2+^ at 20 and 100 μM and progress curves for iron oxidation at A_315_ were measured in a UV/visible spectrophotometer. Enzyme free controls for background oxidation of Fe^2+^ are shown for reference. (**C**) Histogram of the iron loading capacity per biological assembly of EncFtn_sH_, encapsulin, EncFtn-Enc and apoferritin. The results shown are for three technical replicates and represent the optimal iron loading by the complexes after three hours when incubated with Fe^2+^.

**Figure 8-supplement 1.**
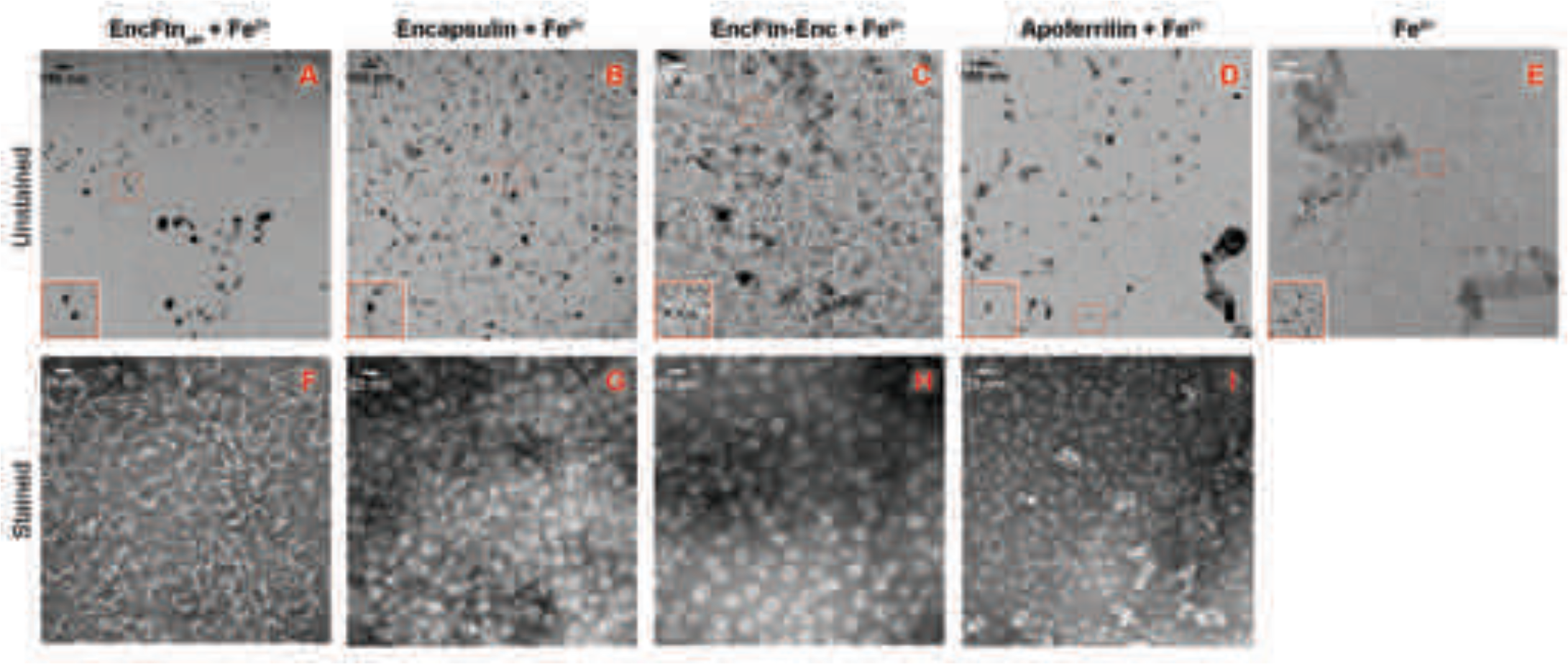
TEM visualization of iron-loaded bacterial nanocompartments and ferritin. Decameric EncFtn_sH_, encapsulin, EncFtn-Enc and apoferritin, at 8.5 μM, were mixed with 147 μM, 1 mM, 1 mM and 215 μM acidic Fe(NH_4_)_2_(SO_4_)_2_, respectively. Protein mixtures were incubated at room temperature for 1 hour prior to TEM analysis with or without uranyl acetate stain. (**A-D**) Unstained EncFtn_sH_, encapsulin, EncFtn-Enc, apoferritin loaded with Fe^2+^, respectively, with 35,000 *x* magnification and scale bars indicate 100 nm. (**E**) Protein-free sample as a control. (**F-I**) Stained EncFtn_sH_, encapsulin, EncFtn-Enc, apoferritin loaded with Fe^2+^, respectively, with 140,000 × magnification and scale bars indicate 25 nm.

**Figure 9.**
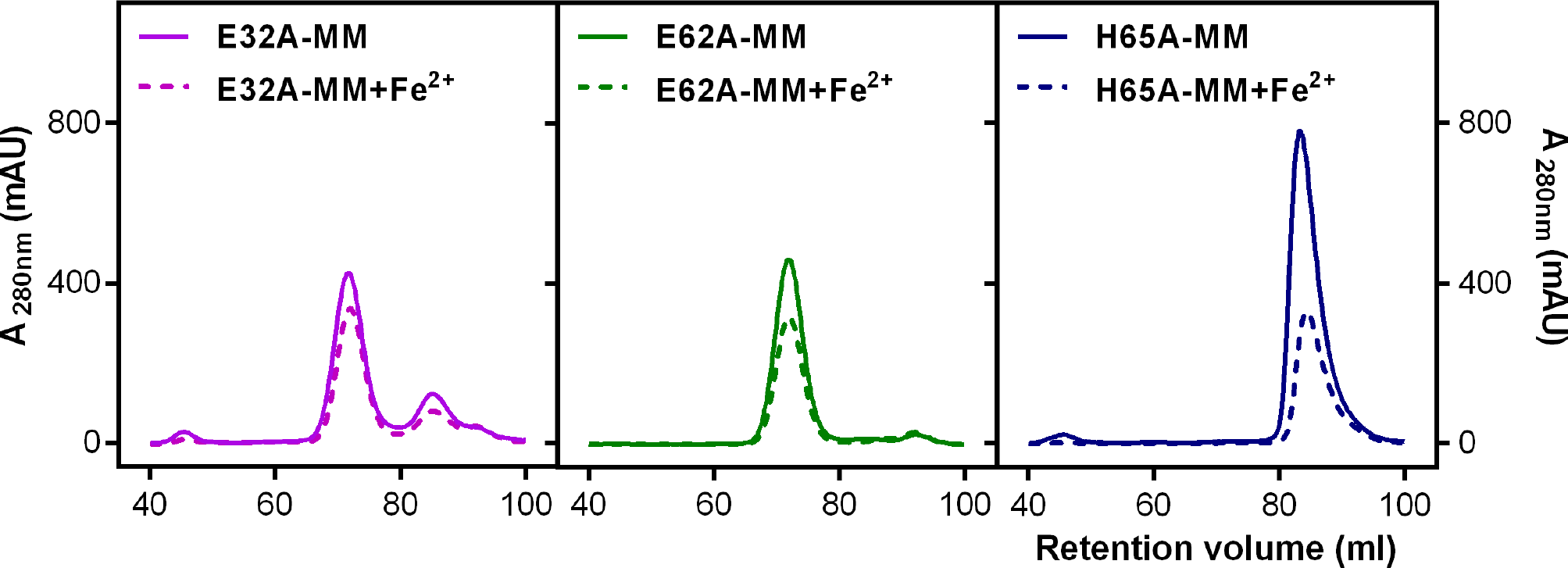
Purification of recombinant *Rhodospirillum rubrum* EncFtn_sH_ FOC mutants. Single mutants E32A, E62A, and H65A of EncFtn_sH_ produced from *E. coli* BL21(DE3) cells grown in minimal medium and minimal medium supplemented with iron were subjected to S200 SEC. (**A**) Gel-filtration chromatogram of the E32A mutant form of EncFtn_sH_ resulted in an elution profile with a majority of the protein eluting as the decameric form of the protein and a small proportion of monomer. (**B**) Gel-filtration of the E62A mutant form of EncFtn_sH_ resulted in an elution profile with a single major decameric peak. (**C**) Gel-filtration of the H65A mutant form of EncFtn_sH_ resulted in a single peak corresponding to the protein monomer.

**Figure 10.**
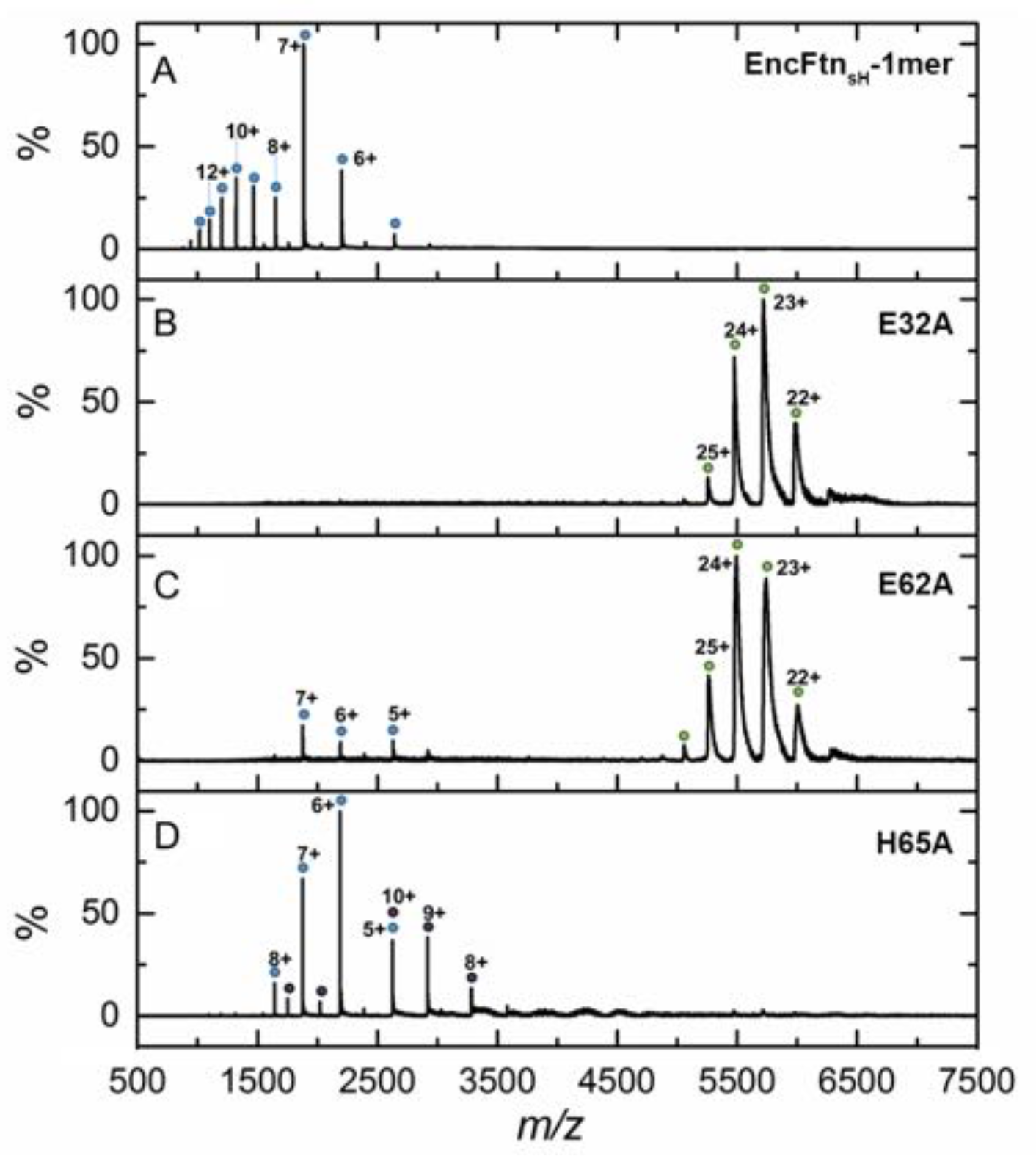
Native mass spectrometry of EncFtn_sH_ mutants. All spectra were acquired in 100 mM ammonium acetate, pH 8.0 with a protein concentration of 5 μM. (A) WT EncFtn_sH_ in the absence of iron displays a charge state distribution consistent with a monomer (see also figure 8). (B) E32A EncFtn_sH_ displays a charge states consistent with a decamer (green circles), a minor species, consistent with the monomer of E32A is also observed (blue circles). (C) E62A EncFtn_sH_ displays a charge states consistent with a decamer (green circles). (D) H65A EncFtn_sH_ displays charge states consistent with both monomer (blue circles) and dimer (purple circles).

**Figure 11.**
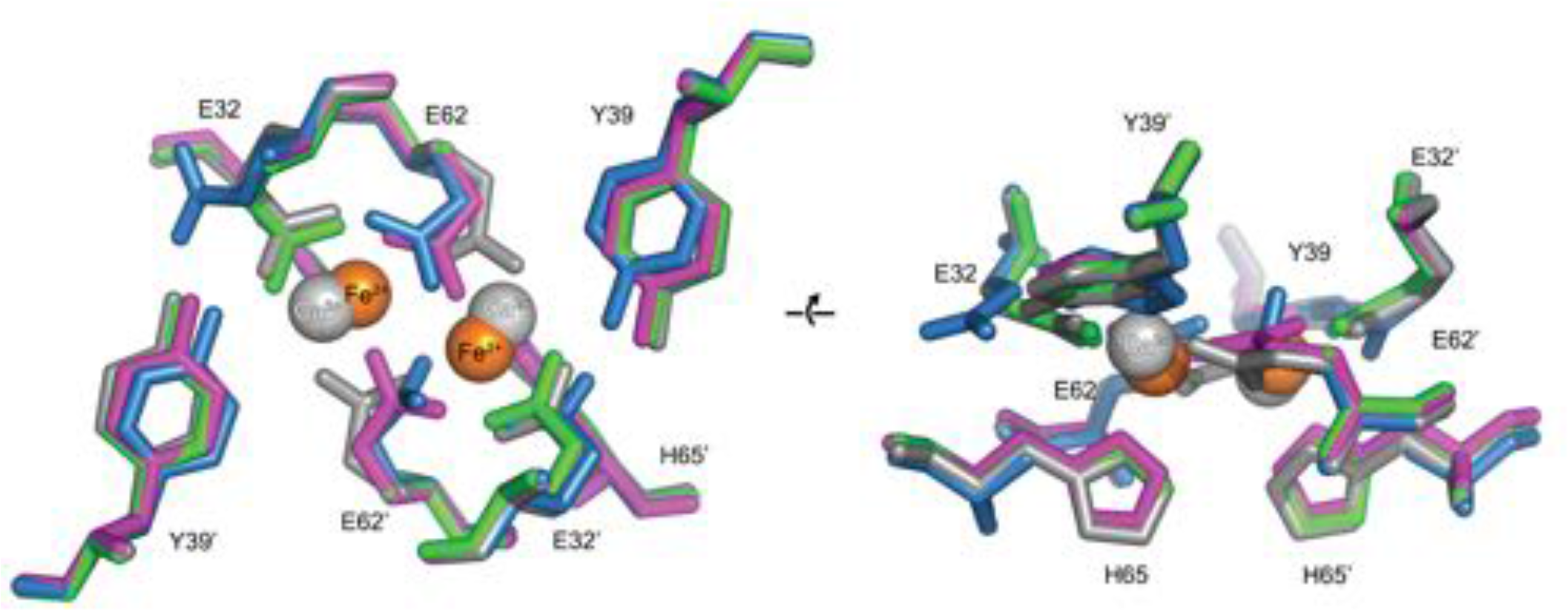
Comparison of the EncFtn_sH_ FOC mutants *vs* wild type. The structures of the three EncFtn_sH_ mutants were all determined by X-ray crystallography. The E32A, E62A and H65A mutants were crystallized in identical conditions to the wild type EncFtn_sH_ structure and were essentially isomorphous in terms of their unit cell dimensions. The FOC residues of the mutants and native EncFtn_sH_ structures are shown as sticks with coordinated Fe^2+^ as orange and Ca^2+^ as grey spheres and are colored as follows: wild type, grey; E32A, pink; E62A, green; H65A, blue. Of the mutants, only H65A has any coordinated metal ions, which appear to be calcium ions from the crystallization condition. The overall organization of FOC residues is retained in the mutants, with almost no backbone movements. Significant differences center around Y39, which moves to coordinate the bound calcium ions in the H65A mutant; and E32, which moves away from the metal ions in this structure.

**Figure 11 supplement 1.**
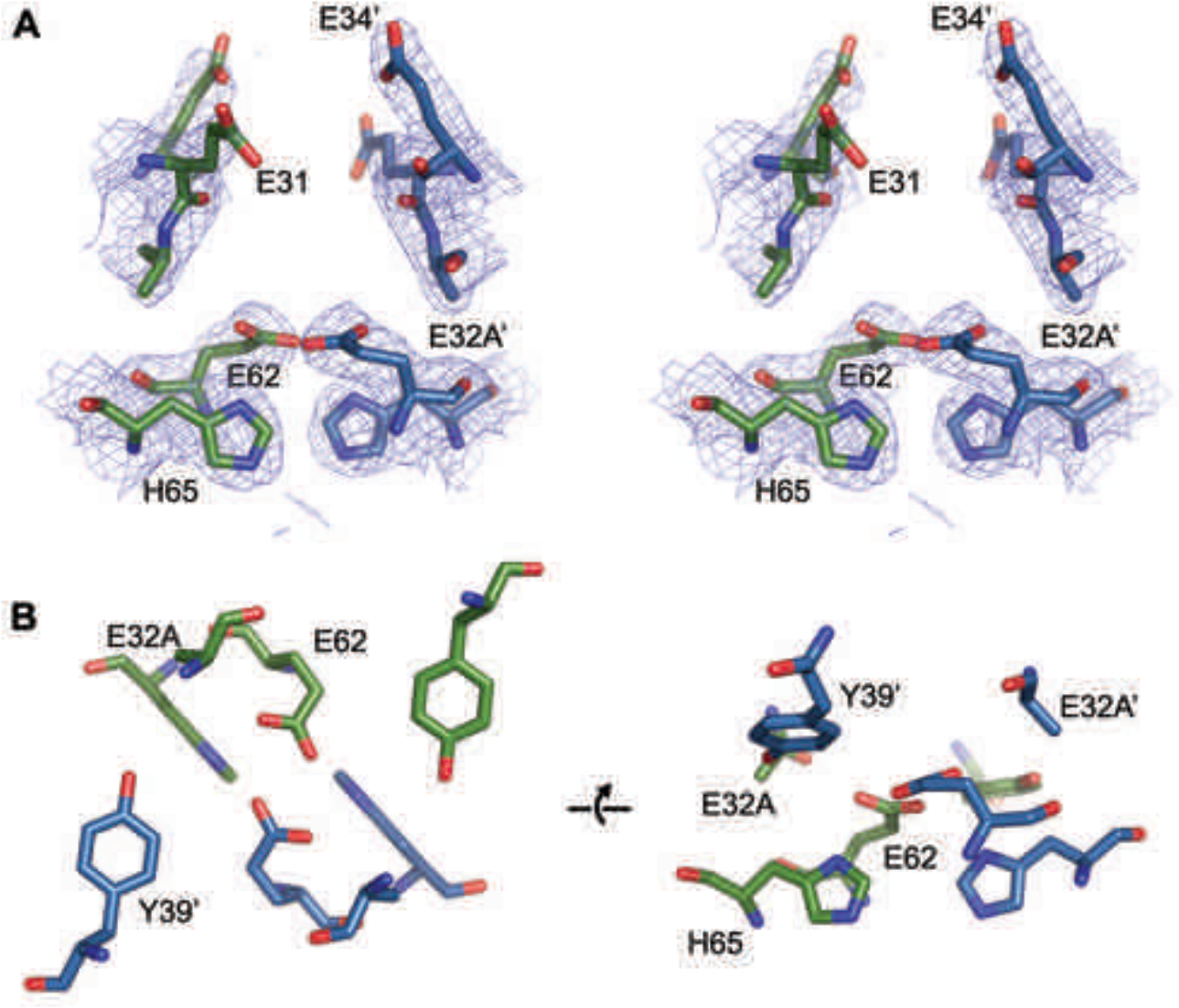
FOC dimer interface of EncFtn_sH_ E32A mutant. (**A**) Wall-eyed stereo view of the metal-binding dimerization interface of EncFtn_sH-E32A_. Protein residues are shown as sticks with blue and green carbons for the different subunits. The 2mFo-DFc electron density map is shown as a blue mesh contoured at 1.5 σ. (**B**) Views of the FOC of the EncFtn_sH-E32A_ mutant. Protein atoms shown as in (**A**).

**Figure 11 supplement 2.**
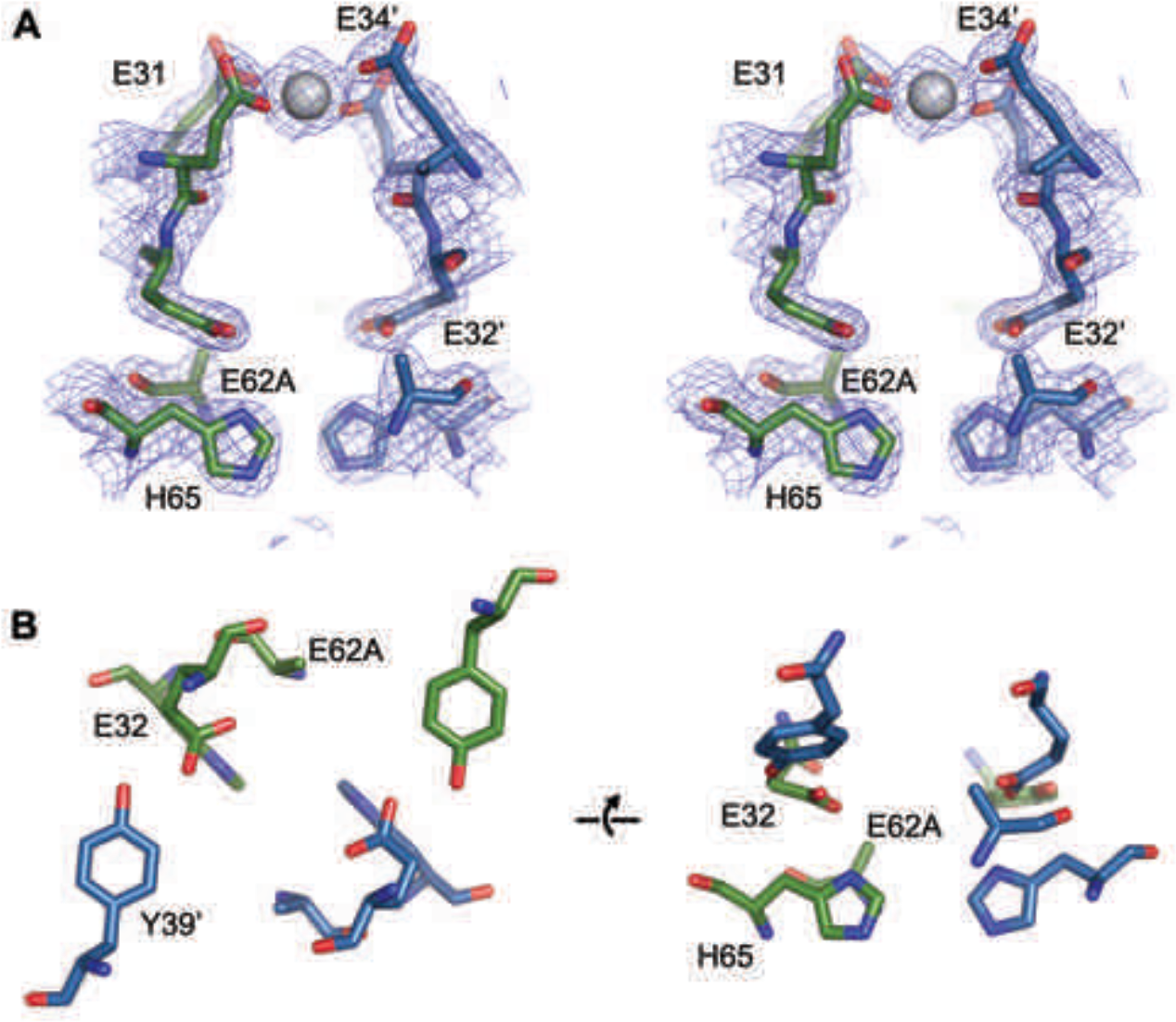
FOC dimer interface of EncFtn_sH_ E62A mutant. (**A**) Wall-eyed stereo view of the metal-binding dimerization interface of EncFtn_sH-E62A_. Protein residues are shown as sticks with blue and green carbons for the different subunits. The 2mFo-DFc electron density map is shown as a blue mesh contoured at 1.5 σ. The single coordinated calcium ion is shown as a grey sphere. (**B**) Views of the FOC of the EncFtn_sH-E62A_ mutant. Protein atoms shown as in (**A**).

**Figure 11 supplement 3.**
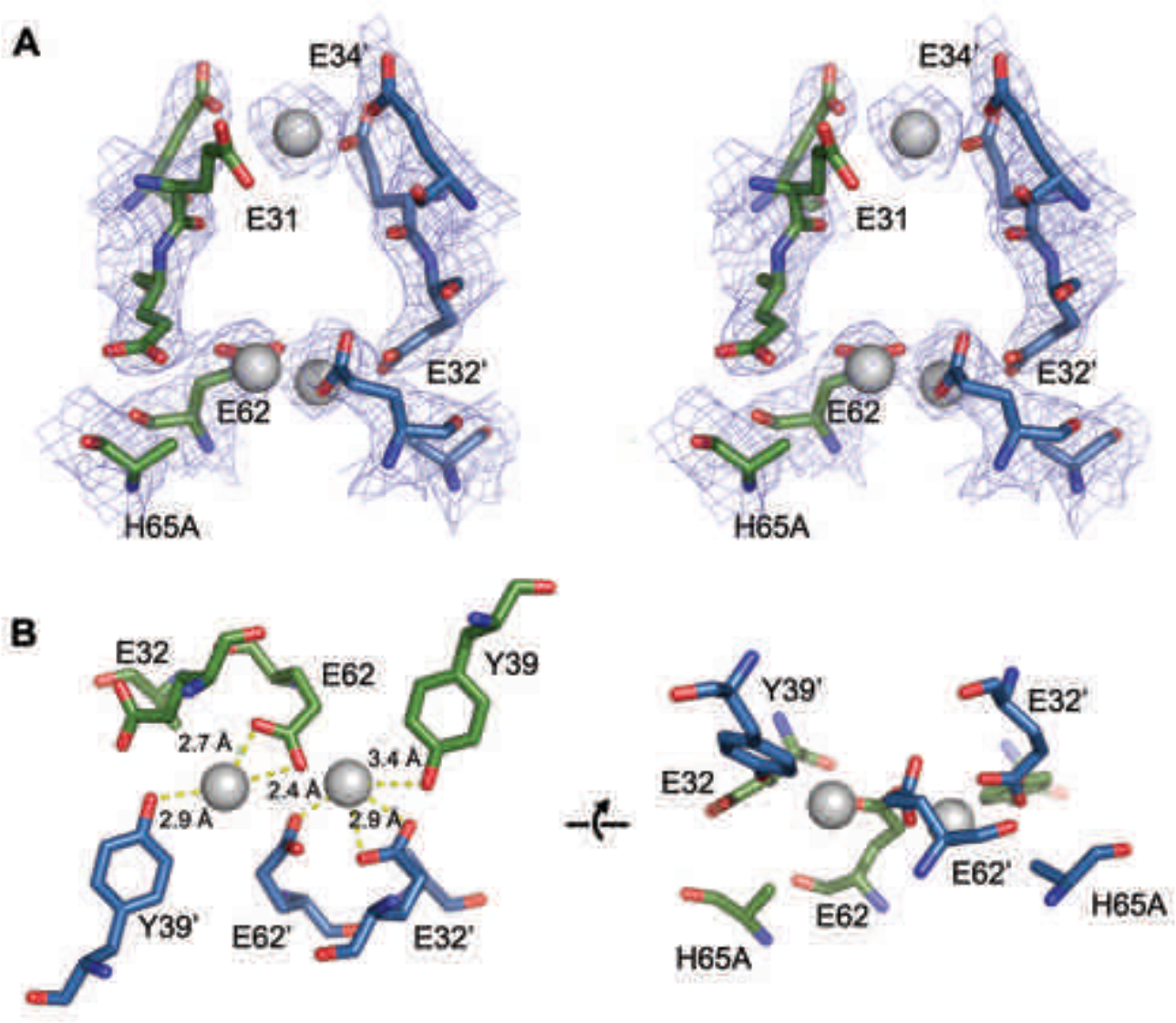
FOC dimer interface of EncFtn_sH_ H65A mutant. (**A**) Wall-eyed stereo view of the metal-binding dimerization interface of EncFtn_sH-H65A_. Protein residues are shown as sticks with blue and green carbons for the different subunits. The 2mFo-DFc electron density map is shown as a blue mesh contoured at 1.5 σ. The coordinated calcium ions are shown as a grey spheres with coordination distances in the FOC highlighted with yellow dashed lines. (**B**) Views of the FOC of the EncFtn_sH-H65A_ mutant. Protein atoms and metal ions shown as in (**A**).

**Figure 12.**
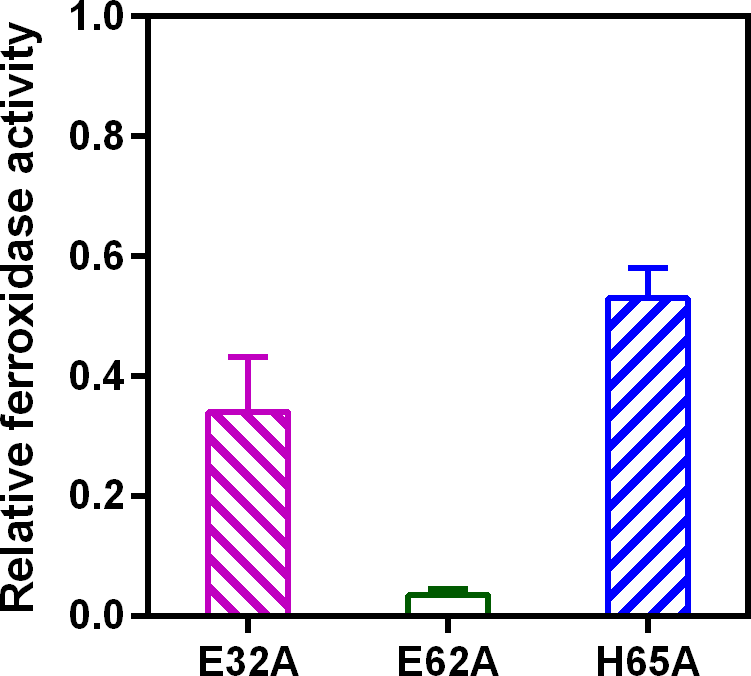
Ferroxidase activity of EncFtn_sH_ mutants. EncFtn_sH_, and the mutant forms E32A, E62A and H65A, each at 20 μM, were mixed with 100 μM acidic Fe(NH_4_)_2_(SO_4_)_2_. Relative ferroxidase activity of the mutant forms is plotted as determined by measuring the absorbance at 315 nm for 1800 s at 25 °C as an indication of Fe^3+^ formation, these data were corrected for background iron oxidation and the final measurements are plotted as a proportion of the activity of the wild-type protein. Three technical repeats were performed and the plotted error bars represent the calculated standard deviations. The FOC mutants showed reduced ferroxidase activity to varied extents, among which E62A significantly abrogated the ferroxidase activity.

**Figure 12-supplement 1.**
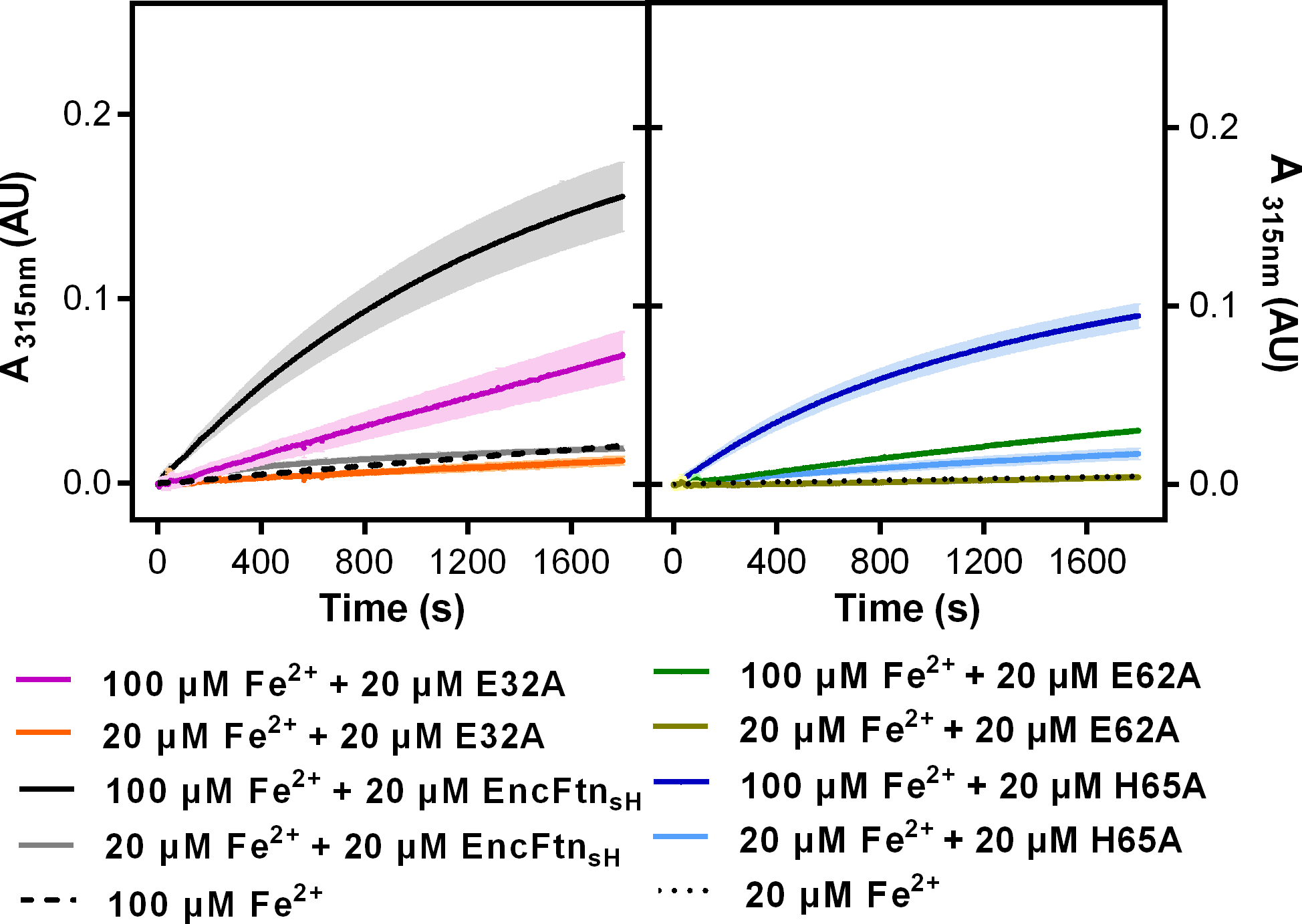
Progress curves recording ferroxidase activity of EncFtn_sH_ mutants. 20 pM EncFtn_sH_, E32A, E62A and H65A were mixed with 20 μM or 100 μM acidic Fe(NH_4_)2(SO_4_)2, respectively. Absorbance at 315 nm was recorded for 1800 s at 25 °C as an indication of Fe^3+^ formation. Protein free samples (dashed and dotted lines) were measured for Fe^2+^ background oxidation as controls. Assays were performed with three technical repeats. Error bars were showed in shadows behind each curves.

**Figure 13.**
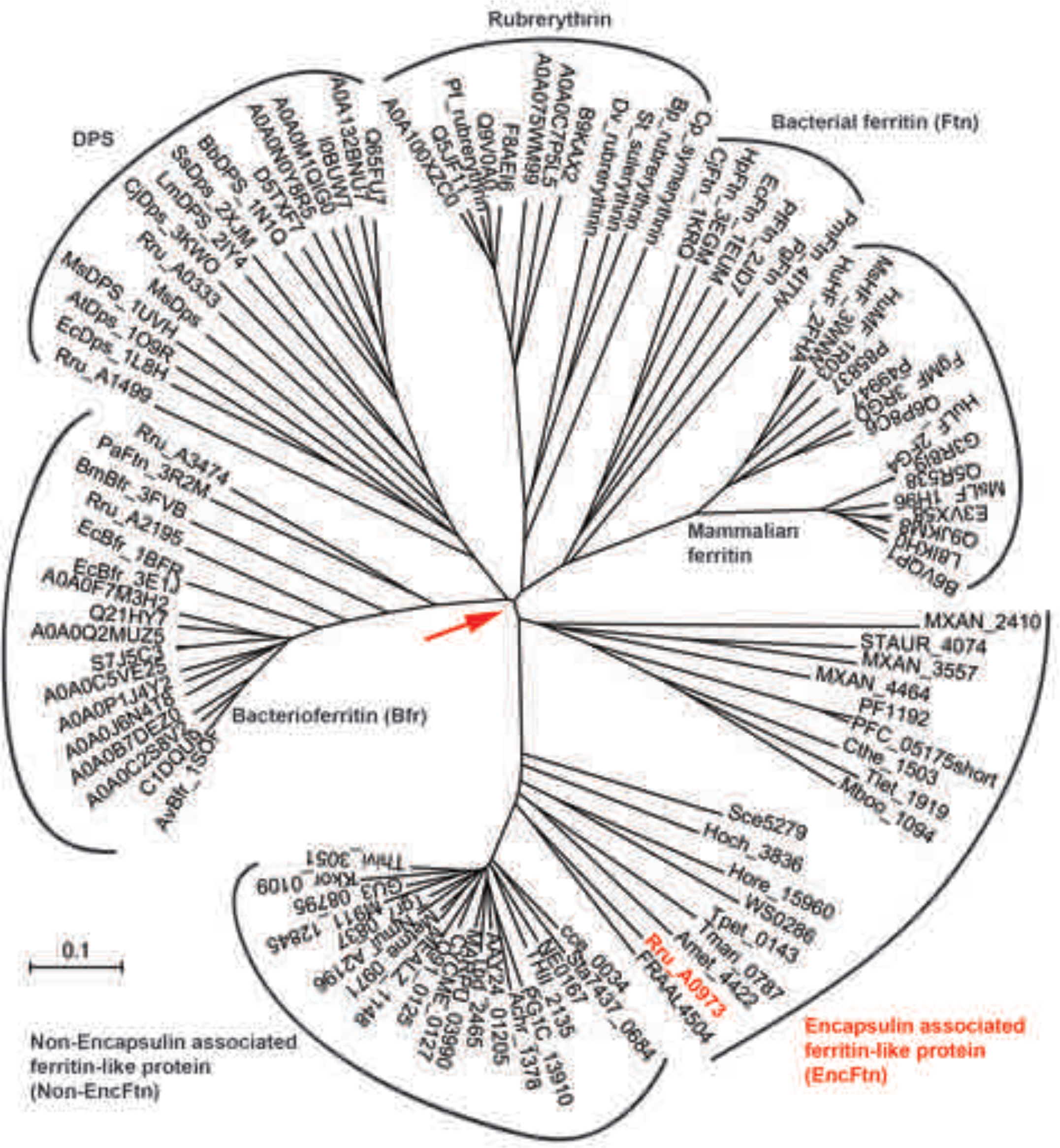
Phylogenetic tree of ferritin family proteins. The tree was built using the Neighbor-Joining method^37^ based on step-wise amino acid sequence alignment of the four-helical bundle portions of ferritin family proteins (Supplementary-file-1). The tree is drawn to scale, with branch lengths in the same units as those of the evolutionary distances used to infer the phylogenetic tree; the likely root of the tree is indicated by a red arrow. The evolutionary distances were computed using the p-distance method^38^ and are in the units of the number of amino acid differences per site. The rate variation among sites was modeled with a gamma distribution (shape parameter = 2.5). The analysis involved 104 amino acid sequences. All ambiguous positions were removed for each sequence pair. There were a total of 262 positions in the final dataset. Evolutionary analyses were conducted in MEGA7^39^.

**Figure 14.**
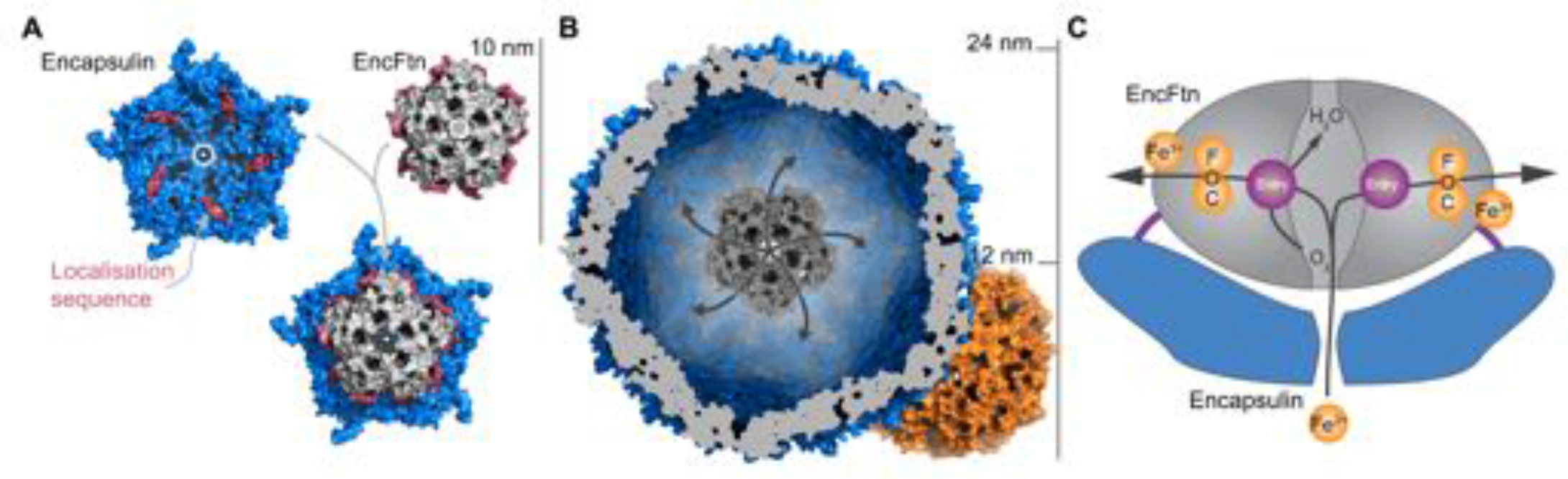
Model of encapsulin nanocompartments. (**A**) Model of EncFtn_sH_ docking to the encapsulin shell. A single pentamer of the icosahedral *T. maritima* encapsulin structure (PDBID: 3DKT)^1^ is shown as a blue surface with the encapsulin localization sequence of EncFtn shown as a purple surface. The C-terminal regions of the EncFtn subunits correspond to the position of the localization sequences seen in 3DKT. Alignment of EncFtn_sH_ with 3DKT positions the central channel directly above the pore in the 3DKT pentamer axis (shown as a grey pentagon). (**B**) Surface view of EncFtn within the encapsulin nanocompartment (grey and blue respectively). The lumen of the encapsulin nanocompartment is considerably larger than the interior of ferritin (shown in orange behind the encapsulin for reference) and thus allows the storage of significantly more iron. The proposed pathway for iron movement through the encapsulin shell and EncFtn FOC is shown with arrows. (**C**) Model of mineralization by EncFtn. As EncFtn is unable to mineralize iron on its surface directly, Fe^2+^ must pass through the encapsulin shell to access the first metal binding site within the central channel of EncFtn_sH_ (entry site) prior to oxidation within the FOC and release as Fe^3+^ to the outer surface of the protein where it can be mineralized within the lumen of the encapsulin cage.

**Table 1.**
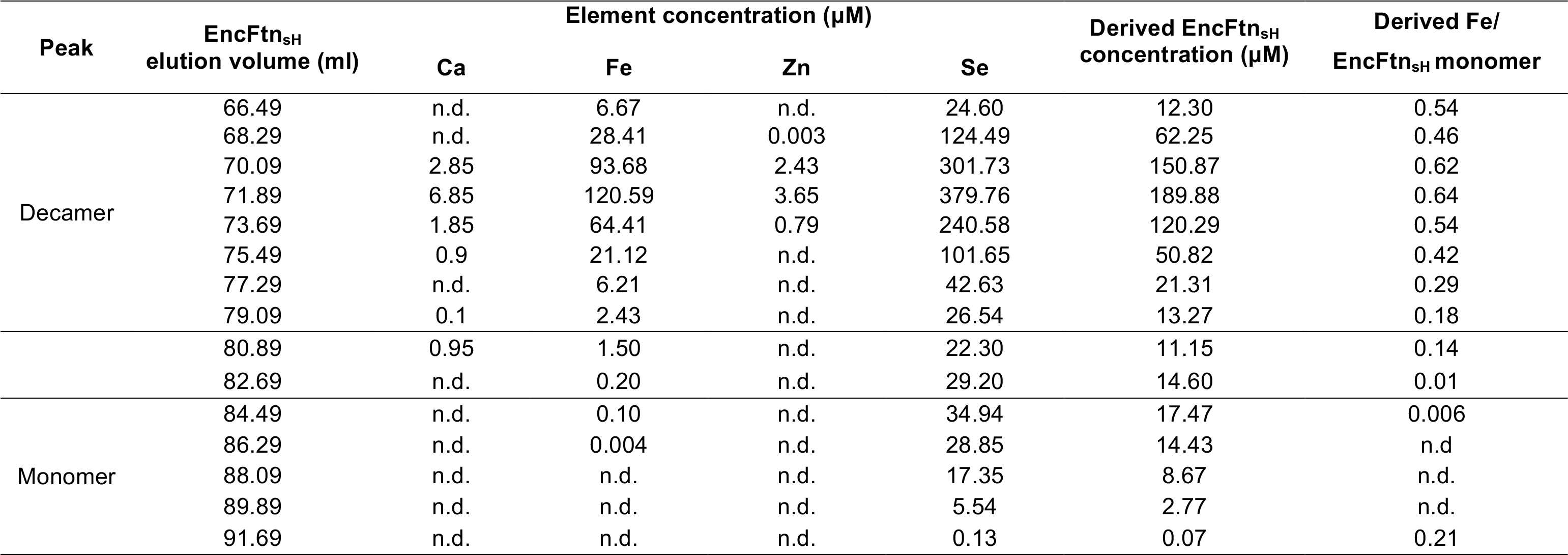
ICP-MS data. EncFtn_sH_ was purified as a SeMet derivative from *E. coli* B834(DE3) cells grown in minimal media with amino acids and supplemented with 1 mM Fe(NH_4_)_2_(SO_4_)_2_. Fractions from size-exclusion gel filtration were collected, acidified and analysed by ICP-MS. EncFtn_sH_ concentration was calculated based on the presence of two SeMet per mature monomer. Samples where the element was undetectable are labelled with n.d‥

**Table 2.** Estimates of EncFtn_sH_ molecular weight from SEC-MALLS analysis. EncFtn_sH_ was purified from *E. coli* BL21(DE3) grown in minimal media by nickel affinity and size exclusion gel filtration chromatography. Fractions from two peaks (decamer and monomer) were pooled separately (Figure 1c) and analysed by SEC-MALLS using a superdex-200 10/300 GL column and Malvern Instruments Viscotek SEC-MALLS instrument (Figure 2C). The decamer and monomer peaks were both symmetric and monodisperse, allowing the estimation of the molecular weight of the species in these fractions^40^. The molecular weights are quoted to the nearest kDa due to the resolution limit of the instrument.

| Molecular Weight (kDa) | Decamer peak | Monomer peak |
| --- | --- | --- |
| Theoretical | 133 | 13 |
| EncFtn<sub>sH</sub>-decamer fractions | 132 | 15 |
| EncFtn<sub>sH</sub>-monomer fractions | 126 | 13 |

**Table 3.** Gel-filtration peak area ratios for EncFtn_sH_ decamer and monomer on addition of different metal ions. EncFtn_sH_ was produced in *E. coli* BL21 (DE3) cultured in minimal media (**MM**) and MM with 1 mM Fe(NH_4_)2(SO_4_)2 (MM+Fe^2+^) and purified by gel-filtration chromatography using an S200 column (GE Healthcare). Monomer fractions of EncFtn_sH_ purified from MM were pooled and run in subsequent analytical gel-filtration runs over the course of three days. Samples of EncFtn_sH_ monomer were incubated with one molar equivalent of metal ion salts at room temperature for two hours before analysis by analytical size-exclusion gel filtration using a Superdex 10/300 GL column. The area for resulting protein peaks were calculated using the Unicorn software (GE Healthcare); peak ratios were calculated to quantify the propensity of EncFtn_sH_ to multimerize in the presence of the different metal ions. The change in monomer:decamer ratios over the three days of experiments may be a consequence of experimental variability, or the propensity of this protein to equilibrate towards decamer over time. The increased decamer:monomer ratio seen in the presence of Fe^2+^, Co^2+^, and Zn^2+^ indicates that these metal ions facilitate multimerization of the EncFtn_sH_ protein, while the other metal ions tested do not appear to induce multimerization.

| Method | Sample | Monomer area | Decamer area | Decamer/Monomer |
| --- | --- | --- | --- | --- |
| Gel filtration S200 chromatography | EncFtn<sub>SH</sub>-MM | 64.27 | 583.58 | 0.11 |
|  | EncFtn<sub>SH</sub>-MM+Fe<sup>2+</sup> | 1938.35 | 426.36 | 4.54 |
| Analytical Gel filtration Day1 | EncFtn<sub>SH</sub>-decamer fractions | 20.24 | 1.80 | 11.24 |
|  | **EncFtn<sub>SH</sub>-monomer fractions** | **2.88** | **21.90** | **0.13** |
|  | Fe(NH<sub>4</sub>)<sub>2</sub>(SO<sub>4</sub>)<sub>2</sub>/ EncFtn<sub>SH</sub>-monomer | 10.99 | 13.03 | 0.84 |
|  | FeSO<sub>4</sub>-HCl/EncFtn<sub>SH</sub>-monomer | 11.28 | 11.37 | 0.99 |
| Analytical Gel filtration Day2 | **EncFtn<sub>SH</sub>-monomer fractions** | **8.30** | **22.76** | **0.36** |
|  | CoCl<sub>2</sub>/EncFtn<sub>SH</sub>-monomer | 17.68 | 14.47 | 1.22 |
|  | MnCl<sub>2</sub>/EncFtn<sub>SH</sub>-monomer | 3.10 | 30.53 | 0.10 |
|  | ZnSO<sub>4</sub>/EncFtn<sub>SH</sub>-monomer | 20.42 | 9.03 | 2.26 |
|  | FeCl<sub>3</sub>/EncFtn<sub>SH</sub>-monomer | 3.87 | 28.57 | 0.14 |
| Analytical Gel filtration Day3 | **EncFtn<sub>SH</sub>-monomer fractions** | **6.31** | **23.41** | **0.27** |
|  | MgSO<sub>4</sub>/EncFtn<sub>SH</sub>-monomer | 5.75 | 30.15 | 0.19 |
|  | Ca acetate/EncFtn<sub>SH</sub>-monomer | 5.54 | 25.18 | 0.22 |

**Table 4.** Data collection and refinement statistics. Statistics for the highest-resolution shell are shown in parentheses. Friedel mates were averaged when calculating reflection numbers and statistics.

|  | WT | E32A | E62A | H65A |
| --- | --- | --- | --- | --- |
| **Data collection** |  |  |  |  |
| Wavelength (Å) | 1.739 | 1.731 | 1.729 | 1.741 |
| Resolution range (Å) | 49.63 - 2.06<br>(2.10 - 2.06) | 48.84 - 2.59 (2.683 -<br>2.59) | 48.87 - 2.21<br>(2.29 - 2.21) | 48.86 - 2.97<br>(3.08 - 2.97) |
| Space group | *P* 1 2<sub>1</sub> 1 | *P* 1 2<sub>1</sub> 1 | *P* 1 2<sub>1</sub> 1 | *P* 1 2<sub>1</sub> 1 |
| Unit cell (Å) *a* | 98.18 | 97.78 | 98.09 | 98.03 |
| *b* | 120.53 | 120.28 | 120.23 | 120.29 |
| *c* | 140.3 | 140.53 | 140.36 | 140.43 |
| β (°) | 95.36 | 95.41 | 95.50 | 95.39 |
| Total reflections | 1,264,922<br>(41,360) | 405,488<br>(36,186) | 1,069,345<br>(95,716) | 323,853<br>(32,120) |
| Unique reflections | 197,873<br>(8,766) | 100,067<br>(9,735) | 162,379<br>(15,817) | 66,658<br>(6,553) |
| Multiplicity | 6.4 (4.7) | 4.1 (3.7) | 6.6 (6.1) | 4.9 (4.9) |
| Anomalous multiplicity | 3.2 (2.6) | N/A | N/A | N/A |
| Completeness (%) | 99.2 (88.6) | 0.99 (0.97) | 1.00 (0.97) | 1.00 (0.99) |
| Anomalous completeness (%) | 96.7 (77.2) | N/A | N/A | N/A |
| Mean I/sigma(I) | 10.6 (1.6) | 8.46 (1.79) | 13.74 (1.80) | 8.09 (1.74) |
| Wilson B-factor | 26.98 | 40.10 | 33.97 | 52.20 |
| R<sub>merge</sub> | 0.123 (0.790) | 0.171 (0.792) | 0.0979 (1.009) | 0.177 (0.863) |
| R<sub>meas</sub> | 0.147 (0.973) | 0.196 (0.923) | 0.1064 (1.107) | 0.199 (0.966) |
| CC1/2 | 0.995 (0.469) | 0.985 (0.557) | 0.998 (0.642) | 0.989 (0.627) |
| CC* | 0.999 (0.846) | 0.996 (0.846) | 0.999 (0.884) | 0.997 (0.878) |
| Image DOI | [10.7488/ds/1342](https://dx.doi.org/10.7488/ds/1342) | [10.7488/ds/1419](https://dx.doi.org/10.7488/ds/1419) | [10.7488/ds/1420](https://dx.doi.org/10.7488/ds/1420) | [10.7488/ds/1421](https://dx.doi.org/10.7488/ds/1421) |
| **Refinement** |  |  |  |  |
| R<sub>work</sub> | 0.1712 (0.3176) | 0.183 (0.288) | 0.165 (0.299) | 0.186 (0.273) |
| R<sub>free</sub> | 0.2061 (0.3445) | 0.225 (0.351) | 0.216 (0.364) | 0.237 (0.325) |
| Number of non-hydrogen atoms | 23,222 | 22,366 | 22,691 | 22,145 |
| macromolecules | 22,276 | 22,019 | 21,965 | 22,066 |
| ligands | 138 | 8 | 24 | 74 |
| water | 808 | 339 | 702 | 5 |
| Protein residues | 2,703 | 2,686 | 2,675 | 2,700 |
| RMS(bonds) (Å) | 0.012 | 0.005 | 0.011 | 0.002 |
| RMS(angles) (°) | 1.26 | 0.58 | 1.02 | 0.40 |
| Ramachandran favored (%) | 100 | 99 | 100 | 99 |
| Ramachandran allowed (%) | 0 | 1 | 0 | 1 |
| Ramachandran outliers (%) | 0 | 0 | 0 | 0 |
| Clash score | 1.42 | 1.42 | 1.79 | 0.97 |
| Average B-factor (Å<sup>2</sup>) | 33.90 | 42.31 | 41.34 | 47.68 |
| macromolecules | 33.80 | 42.35 | 41.31 | 47.6 |
| ligands | 40.40 | 72.80 | 65.55 | 72.34 |
| solvent | 36.20 | 38.95 | 41.46 | 33.85 |
| PDB ID | [5DA5](https://www.rcsb.org/structure/5DA5) | [5L89](https://www.rcsb.org/structure/5L89) | [5L8B](https://www.rcsb.org/structure/5L8B) | [5L8G](https://www.rcsb.org/structure/5L8G) |

**Table 5.**
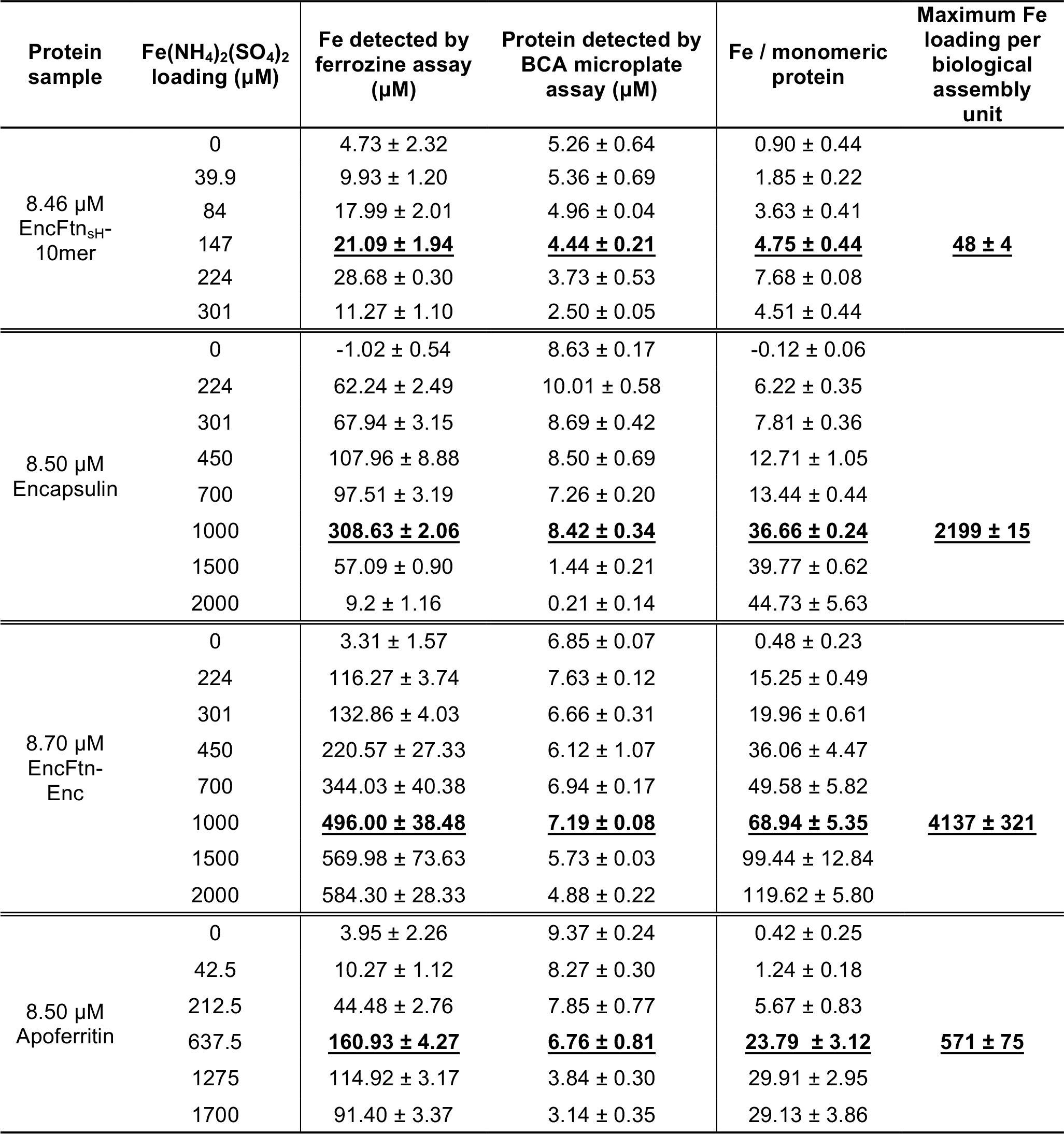
Iron loading capacity of EncFtn, encapsulin and ferritin. Protein samples (at 8.5 μM) including decameric EncFtn_sH_, Encapsulin, EncFtn-Enc and apoferritin were mixed with Fe(NH_4_)_2_(SO_4_) (in 0.1%(v/v) HCl) of different concentrations in 50 mM Tris-HCl (pH 8.0), 150 mM NaCl buffer at room temperature for 3 hours in the air. Protein-Fe mixtures were centrifuged at 13,000 *x* g to remove precipitated material and desalted prior to the Fe and protein content analysis by ferrozine assay and BCA microplate assay, respectively. Fe to protein ratio was calculated to indicate the Fe binding capacity of the protein. Protein stability was compromised at high iron concentrations; therefore, the highest iron loading with the least protein precipitation was used to derive the maximum iron loading capacity per biological assembly (Underlined and highlighted in bold). The biological unit assemblies are a decamer for EncFtn_sH_, a 60mer for encapsulin, a 60mer of encapsulin loaded with 12 copies of decameric EncFtn in the complex, and 24mer for horse spleen apoferritin. Errors are quoted as the standard deviation of three technical repeats in both the ferrozine and BCA microplate assays.

## Methods

### Cloning

Genes of interest were amplified by PCR using *R. rubrum* ATCC 11170 genomic DNA (DSMZ) as the template and KOD Hot Start DNA Polymerase (Novagen). Primers used in this study are listed in Supplementary file 2. PCR products were visualized in 0.8% agarose gel stained with SYBR Safe (Life Technologies). Fragments of interest were purified by gel extraction (Qiagen) before digestion by endonuclease restriction enzymes (Thermo Fisher Scientific) at 37 °C for 1 hour, followed by ligation with similarly digested vector pET-28a(+) or pACYCDuet-1 at room temperature for 1 hour. Ligation product was transformed into chemically competent *Escherichia coli* Top10 cells and screened against 50 ng/μl kanamycin for pET-28a(+) based constructs or 34 ng/μl chloramphenicol for pACYCDuet-1 based constructs. DNA insertion was confirmed through Sanger sequencing (Edinburgh Genomics). Sequence verified constructs were transformed into *E. coli* BL21(DE3) or Tuner(DE3) for protein production. Alternatively, plasmids transformed into *E. coli* B834(DE3) cells were cultured in selenomethionine media.

### Protein production and purification

A single colony of *E. coli* BL21(DE3) or Tuner(DE3) cells, transformed with protein expression plasmid, was transferred into 10 ml LB medium, or M9 minimal medium (MM), supplemented with appropriate antibiotic, and incubated overnight at 37 °C with 200 rpm shaking. The overnight preculture was then inoculated into 1 liter of LB medium and incubated at 37 °C with 200 rpm shaking. Recombinant protein production was induced at OD_600_= 0.6 by the addition of 1 mM IPTG and the incubation temperature was reduced to 18 °C for overnight incubation. Cells were pelleted by centrifugation at 4000 *g* for 20 min at 4 °C, and resuspended 10-fold (volume per gram of cell pellet) in PBS to wash cells before a second centrifugation step. Cells were resuspended in 10-times (v/w) of appropriate lysis buffer for the purification method used (see details of buffers below) and lysed by sonication on ice, with ten cycles of 30-second burst of sonication at 10 μm amplitude and 30 seconds of cooling. Cell lysate was clarified by centrifugation at 20,000 × *g*, 30 minutes, 4 °C; followed by filtration using a 0.22 μm syringe filter (Millipore).

Selenomethionine labelled protein was produced by growing a single colony of *E. coli* B834 (DE3) cells transformed with protein expression plasmids in 100 ml LB medium supplemented with appropriate antibiotic overnight at 37 °C with shaking at 200 rpm. The overnight pre-culture was pelleted by centrifugation 3,000 × *g*, 4 °C, 15 minutes and washed twice with M9 minimal medium. The washed cells were transferred to 1 liter of M9 minimal medium supplemented with nineteen_L_-amino acids (without methionine) plus selenomethionine, 2 mM MgSO_4_, 0.4% (w/v) glucose, 1 mM Fe(NH_4_)_2_(SO_4_)_2_. Cells were incubated at 37 °C with 200 rpm shaking and recombinant protein production was induced at OD_600_= 0.6 by the addition of 1 mM IPTG and the incubation temperature was reduced to 18 °C for overnight incubation. Cells were harvested and lysed as above.

### His-tagged protein purification

Clarified cell lysate was loaded onto a 5 ml HisTrap column (GE Healthcare) pre-equilibrated with HisA buffer (50 mM Tris-HCl, 500 mM NaCl and 50 mM imidazole, pH 8.0). Unbound proteins were washed from the column with HisA buffer. His-tagged proteins were then eluted by a step gradient of 50% HisA buffer and 50% HisB buffer (50 mM Tris-HCl, 500 mM NaCl and 500 mM imidazole, pH 8.0). Fractions containing the protein of interest, as determined by 15% acrylamide (w/v) SDS-PAGE, were pooled before loading onto a gel-filtration column (HiLoad 16/600 Superdex 200, GE Healthcare) equilibrated with GF buffer (50 mM Tris-HCl, pH 8.0, 150 mM NaCl). Fractions were subjected to 15% SDS-PAGE and those containing the protein of interest were pooled for further analysis.

### Sucrose gradient ultracentrifugation purification

Co-expressed encapsulin and EncFtn (EncFtn-Enc) and encapsulin protein were both purified according to the protocol used by M. Sutter^41^. Briefly, EncFtn-Enc or encapsulin was expressed based on pACYCDuet-1 vector. The *E. coli* cells were growing, induced, harvested and sonicated in a similar way as described above. GF buffer used in this purification contains 50 mM Tris-HCl, pH 8.0, and 150 mM NaCl. To remove RNA contamination, the lysate was supplemented with 50 μg/ml RNase A and rotated at 10 rpm and room temperature for 2 hrs, followed by centrifugation at 34,000 *× g* and 4°C for 20 min and filtering through 0.22 μM syringe filter. Proteins were pelleted through 38% (w/v) sucrose cushion by ultracentrifugation at 100,000 × *g* and 4 °C for 21 hrs. 10% – 50% (w/v) sucrose gradient ultracentrifugation was applied to further separate the proteins at 100,000 × *g* and 4 °C for 17 h. Protein was dialyzed against GF buffer to remove sucrose before being used in chemical assays or TEM.

### Transmission electron microscopy

TEM imaging was performed on purified encapsulin, EncFtn, and EncFtn-Enc and apoferritin. Purified protein at 0.1 mg/ml concentration was spotted on glow-discharged 300 mesh carbon-coated copper grids and excess liquid wicked off with Whatman filter paper. The grids were washed with distilled water and blotted with filter paper three times before staining with 0.2% uranyl acetate, blotting and air-drying. Grids were imaged using a JEM1400 transmission electron microscope and images were collected with a Gatan CCD camera. Images were analyzed using ImageJ (NIH) and size-distribution histograms were plotted using Prism6 (GraphPad software). To observe iron mineral formation by TEM, protein samples at 8.5 pM concentration including EncFtn_sH_, Encapsulin, EncFtn-Enc and apoferritin were supplemented with acidic Fe(NH_4_)_2_(SO_4_)_2_ at their maximum iron loading ratio in room temperature for 1 hr. The mixtures were subjected to TEM analysis with or without uranyl acetate staining.

### Protein crystallization and X-ray data collection

EncFtn_sH_ was purified by anion exchange and S200 size-exclusion gel-filtration and concentrated to 10 mg/ml (based on extinction coefficient calculation). Crystallization drops were set up using the hanging drop vapor diffusion method at 292 K. Glass coverslips were set up with 1-2 μl protein mixed with 1 μl well solution (0.14 M calcium acetate, 15% (w/v) PEG 3350) and sealed over 1 ml of well solution. Crystals appeared after 5 days and were harvested from the well using a LithoLoop (Molecular Dimensions Limited), transferred briefly to a cryoprotection solution containing well solution supplemented with 1 mM FeSO_4_ (in 0.1% (v/v) HCl), 20% (v/v) PEG 200, and subsequently flash cooled in liquid nitrogen. Crystals of the EncFtn_sH_ point mutants were produced in the same manner as for the EncFtn_sH_ wild-type protein.

All crystallographic datasets were collected on the macromolecular crystallography beamlines at Diamond Light Source (Didcot, UK) at 100 K using Pilatus 6M detectors. Diffraction data were integrated and scaled using XDS^42^ and symmetry related reflections were merged with Aimless^43^. Data collection statistics are shown in Table 4. The resolution cut-off used for structure determination and refinement was determined based on the CC_1/2_ criterion proposed by Karplus and Diederichs^44^.

The structure of EncFtn_sH_ was determined by molecular replacement using PDB ID: 3K6C as the search model, modified to match the sequence of the target protein using Chainsaw^45^. A single solution comprising three decamers in the asymmetric unit was found by molecular replacement using Phaser^46^. The initial model was rebuilt using Phenix.autobuild^47^ followed by cycles of refinement with Phenix.refine^48^, with manual rebuilding and model inspection in Coot^49^. The final model was refined with isotropic B-factors, torsional NCS restraints, and with anomalous group refinement. The model was validated using MolProbity^50^. Structural superimpositions were calculated using Coot. Crystallographic figures were generated with PyMOL. Multiple sequence alignment of EncFtn and ferritin family proteins was performed using Clustal Omega^51^ and displayed with Espript 3.0^52^. Model refinement statistics are shown in Table 4. The final models and experimental data are deposited in the PDB and diffraction image files are available at the Edinburgh DataShare repository.

### Horse spleen apoferritin preparation

Horse spleen apoferritin purchased from Sigma Aldrich was dissolved in deaerated MOPS buffer (100 mM MOPS, 100 mM NaCl, 3 g/100 ml Na_2_S_2_O_4_ ^53^ and 0.5 M EDTA, pH 6.5)^53^. Protein was dialyzed against 1 liter MOPS buffer in room temperature for 2 days before buffer exchanging to GF buffer (50 mM Tris-HCl, pH 8.0, 150 mM NaCl) in a vivaspin column with 5 kDa cut-off (Sartorius) for several times. Fe content of apoferritin was detected using ferrozine assay^54^. Protein concentration was determined using Pierce™ Microplate BCA Protein Assay Kit. Apoferritin containing less than 0.5 Fe per 24-mer was used in the ferroxidase assay. Apoferritin used in the Fe loading capacity experiment was prepared in the same way with 5-15 Fe per 24-mer.

### Ferroxidase assay

1 mM and 200 μM Fe(NH_4_)_2_(SO_4_)_2_ stock solutions were prepared in 0.1% (v/v) HCl anaerobically. Protein solutions with 20 μM FOC were diluted from ~10 mg/ml frozen stock in GF buffer (50 mM Tris-HCl, pH 8.0 and 150 mM NaCl) anaerobically. Ferroxidase activity was initiated by adding 450 μl protein to 50 μl of acidic Fe(NH_4_)_2_(SO_4_)_2_ at the final concentration of 100 μM and 20 μM in the air, respectively. The ferroxidase activity was measured by monitoring the Fe^3+^ formation which gives rise to the change of the absorbance at 315 nm^55^. Absorbance at 315 nm was recorded every second over 1800 seconds using a quartz cuvette in a JASCO V-730 UV/VIS spectrophotometer (JASCO Inc.). In recombinantly coexpressed nanocompartments the ratio of EncFtn to Enc was assumed as 2 to 1, assuming each of the twelve pentameric vertices of the icosahedral encapsulin were occupied with decameric EncFtn. The data are presented as the mean of three technical replicates with error bars indicating one standard deviation from the mean.

### Iron loading capacity of ferritins

In order to determine the maximum iron loading capacity, around 8.5 μM proteins including decameric EncFtn_sH_, Encapsulin, EncFtn-Enc and apoferritin were loaded with various amount of acidic Fe(NH_4_)_2_(SO_4_)_2_ ranging from 0 to 1700 μM. Protein mixtures were incubated in room temperature for 3 hrs before desalting in Zebra™ spin desalting columns (7 kDa cut-off, Thermo Fisher Scientific) to remove free iron ions. The protein concentration was determined using Pierce™ Microplate BCA assay kit (Thermo Fisher Scientific). The protein standard curve was plotted according to the manufacturer. The Fe content in the samples was determined using modified ferrozine assay^56^. Briefly speaking, 100 μl protein sample was mixed with 100 μl mixture of equal volume of 1.4 M HCl and 4.5% (w/v) KMnO_4_ and incubated at 60 °C for 2 hrs. 20 μl of the iron-detection reagent (6.5 mM ferrozine, 6.5 mM neocuproine, 2.5 M ammonium acetate, and 1 M ascorbic acid dissolved in H_2_O) was added to the cooled tubes. 30 min later, 200 μl of the solution was transferred into a well of 96-well plate and the absorbance at 562 nm was measured on the plate reader Spectramax M5 (Molecular Devices). The standard curve was plotted using various concentrations of FeCl_3_ (in 10 mM HCl) diluted in the gel-filtration buffer. Three technical repeats were performed for both the ferrozine and microplate BCA assays. Samples analyzed by ICP-MS were prepared in the same way by mixing protein and ferrous ions and desalting.

### Peroxidase assay

The peroxidase activity of EncFtn_sH_ was determined by measuring the oxidation of *ortho-* phenylenediamine (OP) by H_2_O_2_^25^. EncFtn_sH_ decamer and monomeric fractions purified from minimal media were both used in the assay. Ortho-phenylenediamine was prepared as a 92.5 mM stock solution in 50 mM Tris-HCl (pH 8.0). 80, 70, 60, 50, 40, 30, 20 and 10 mM of OP were prepared by diluting the stock solution in the 50 mM Tris-HCl (pH 8.0). 100 μl of each diluted OP was added to a 96-well plate in 3 repeats. 1 μl of 32 μM protein was supplemented into each well to a final concentration of 160 nM, followed by the addition of 2 μl of 30% H_2_O_2_. After 15 minutes shaking in the dark, the reaction was stopped by adding 100 μl of 0.5 M H_2_SO_4_. The peroxidase activity was measured by monitoring the absorbance at 490 nm in the SpectraMax M5 Microplate Reader (Molecular Devices Corporation)^25^.

### ICP-MS analysis

Protein samples were diluted 50-fold into a solution of 2.5% HNO_3_ (Suprapur, Merck) containing 20 μg/L Pt as internal standard. Matrix-matched elemental standards (containing analyte metal concentrations 0 – 1,000 μg/L) were prepared by serial dilution from individual metal standard stocks (VWR) with identical solution compositions, including the internal standard. All standards and samples were analyzed by ICP-MS using a Thermo x-series instrument operating in collision cell mode (using 3.0 ml min^−1^ flow of 8% H_2_ in He as the collision gas). Isotopes ^44^Ca,^56^Fe, ^66^Zn,^78^Se, and ^195^Pt were monitored using the peak-jump method (100 sweeps, 25-30 ms dwell time on 5 channels per isotope, separated by 0.02 atomic mass units) in triplicate.

### Mass spectrometry analysis

For native MS analysis, all protein samples were buffer exchanged into 100 mM ammonium acetate (pH 8.0; adjusted with dropwise addition of 1% ammonia solution) using Micro Biospin^®^ Chromatography Columns (Bio-Rad, UK) prior to analysis and the resulting protein samples were analyzed at a final concentration of ~5 μM (oligomer concentration). In order to obtain Fe-bound EncFtn, 100 μM or 300 μM of freshly prepared FeCl_2_ was added to apo-EncFtn_sH_ (monomer peak) immediately prior to buffer exchange into 100 mM ammonium acetate (pH 8.0). Samples were analyzed on a quadrupole ion-mobility time of flight instrument (Synapt G2, Waters Corp., Manchester, UK), equipped with a nanomate nanoelectrospray infusion robot (Advion Biosciences). Instrument parameters were tuned to preserve non-covalent protein complexes. After optimization, typical parameters were: nanoelectrospray voltage 1.54 kV; sample cone 50 V; extractor cone 0 V; trap collision voltage 4 V; source temperature 80 °C; and source backing pressure 5.5 mbar. For improved mass resolution the sample cone was raised to 155 V. Ion mobility mass spectrometry (IM-MS) was performed using the travelling-wave mobility cell in the Synapt G2, employing nitrogen as the drift gas. Typically, the IMS wave velocity was set to 300 m/s; wave height to 15 V; and the IMS pressure was 1.8 mbar. For collision cross section determination, IM-MS data was calibrated using denatured equine myoglobin and data was analyzed using Driftscope v2.5 and MassLynx v4.1 (Waters Corp., UK). Theoretical collision cross sections (CCS) were calculated from pdb files using IMPACT software v. 0.9.1^57^. In order to obtain information on the topology of the EncFtn_sH_ assembly, gas-phase dissociation of the Fe-associated EncFtn_sH_ complex was achieved by increasing the sample cone and/or trap collision voltage prior to MS analysis.

### SEC-MALLS

Size-exclusion chromatography (ÅKTA-Micro; GE Healthcare) coupled to UV, static light scattering and refractive index detection (Viscotec SEC-MALS 20 and Viscotec RI Detector:VE3580; Malvern Instruments) were used to determine the molecular mass of fractions decamer and monomer of EncFtn_sH_ in solution individually. Protein concentration was determined by measurement of absorbance at 280 nm and calculated using the extinction coefficient ɛ^0.1%^= 1.462 mg^−1^ ml cm^−1^. 100 μl of 1.43 mg/ml fractions of EncFtn_sH_ decamer and 4.03 mg/ml fractions of EncFtn_sH_ monomer were run individually on a Superdex 200 10/300 GL size exclusion column pre-equilibrated in 50 mM Tris-HCl (pH 8.0), 150 mM NaCl at 22 °C with a flow rate of 0.5 ml/min. Light scattering, refractive index (RI) and A_280nm_ were analyzed by a homo-polymer model (OmniSEC software, v 5.1; Malvern Instruments) using the following parameters for fractions of decamer and monomer: the extinction coefficient (dA / dc) at 280 nm was 1.46 AU mg ml^−1^ and specific refractive index increment (dn / dc) was 0.185 ml g^−1^.

## Metal binding analysis by PAGE

Recombinant EncFtn_sH_ fractions at 50 μM concentration were incubated with one molar equivalent of metal ions at room temperature for 2 hours. Half of each sample was mixed with 5 × native loading buffer (65 mM Tris-HCl, pH 8.5, 20% glycerol and 0.01% bromophenol blue) and run on non-denaturing PAGE gels (10% acrylamide) and run in Tris/glycine buffer, 200 V, 4 °C for 50 min. The remaining samples were left for an additional three hours prior to SDS-PAGE (15% acrylamide) analysis. SDS-PAGE gels were run at room temperature at 200 V, room temperature for 50 min. Gels were stained with Coomassie Brilliant Blue R250 and scanned after de-staining in water.

### Accession codes

Coordinates and structure factors for the structures presented in this paper have been deposited in the PDB under the following accession codes: EncFtn_sH_, 5DA5; EncFtn_sH_-E32A, 5L89; EncFtn_sH_-E62A, 5L8B; EncFtn_sH_-H65A, 5L8G.

## Acknowledgements

This work was supported by a Royal Society Research Grant awarded to JMW (RG130585), a BBSRC New Investigator Grant to JMW and DJC (BB/N005570/1) and a Wellcome Trust Institute Strategic Support Fund grant awarded to set up the XtalPod crystallization facility at the University of Edinburgh. DJC, SH, and KA are funded by the University of Edinburgh. DH is funded by the China Scholarship Council. KJW and ET are funded by the Wellcome Trust and Royal Society through a Sir Henry Dale Fellowship awarded to KJW (098375/Z/12/Z).

We would like to thank the staff on the Macromolecular Crystallography beamlines at Diamond Light Source for their assistance with data collection. We would like to thank Dr Steve Mitchell in the University of Edinburgh and Electron Microscopy Research Services in Newcastle University for the use of Transmission Electron Microscopes. We would like to thank Dr Michael Capeness and Dr. Louise Horsfall for the use of their anaerobic chamber. We would like to thank staff in the Edinburgh Protein Production Facility for their guidance and patience, Dr Janice Brahman, Dr Liz Blackburn, Dr Matthew Nowicki and Dr Martin Wear. We would like to thank Professor Rick Lewis and Dr Arnaud Basle (Newcastle University, UK), and Dr John Berrisford (PDBe, EMBL-EBI, Hinxton, Cambridge) for help identifying the glycolic acid ligand in the ferroxidase center. We would like to thank Dr Dominic Campopiano, Dr Elisabeth Lowe and Laura Tuck for their critical reading of this manuscript and helpful discussions.

---

**Supplementary file 1.**
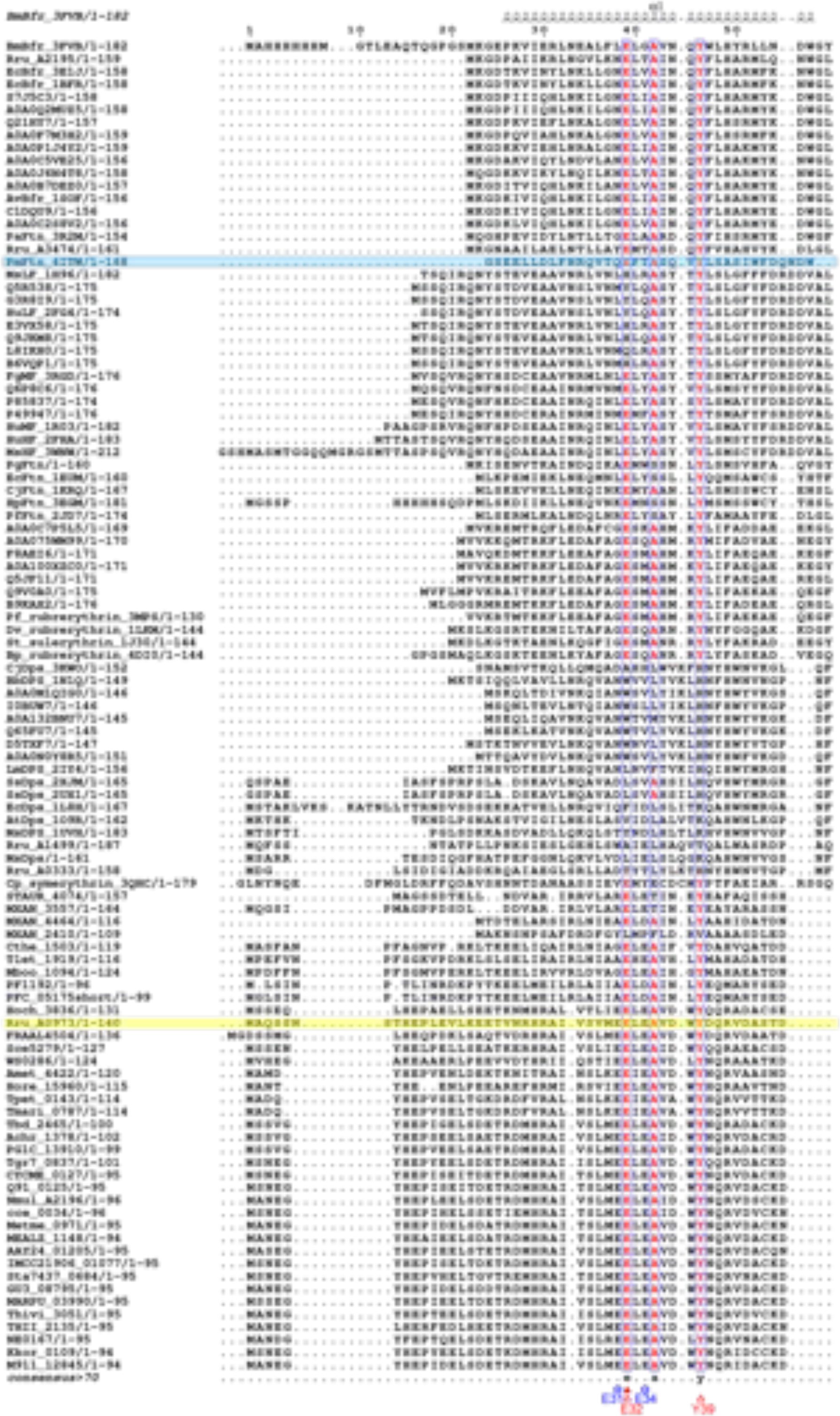
Multiple sequence alignment

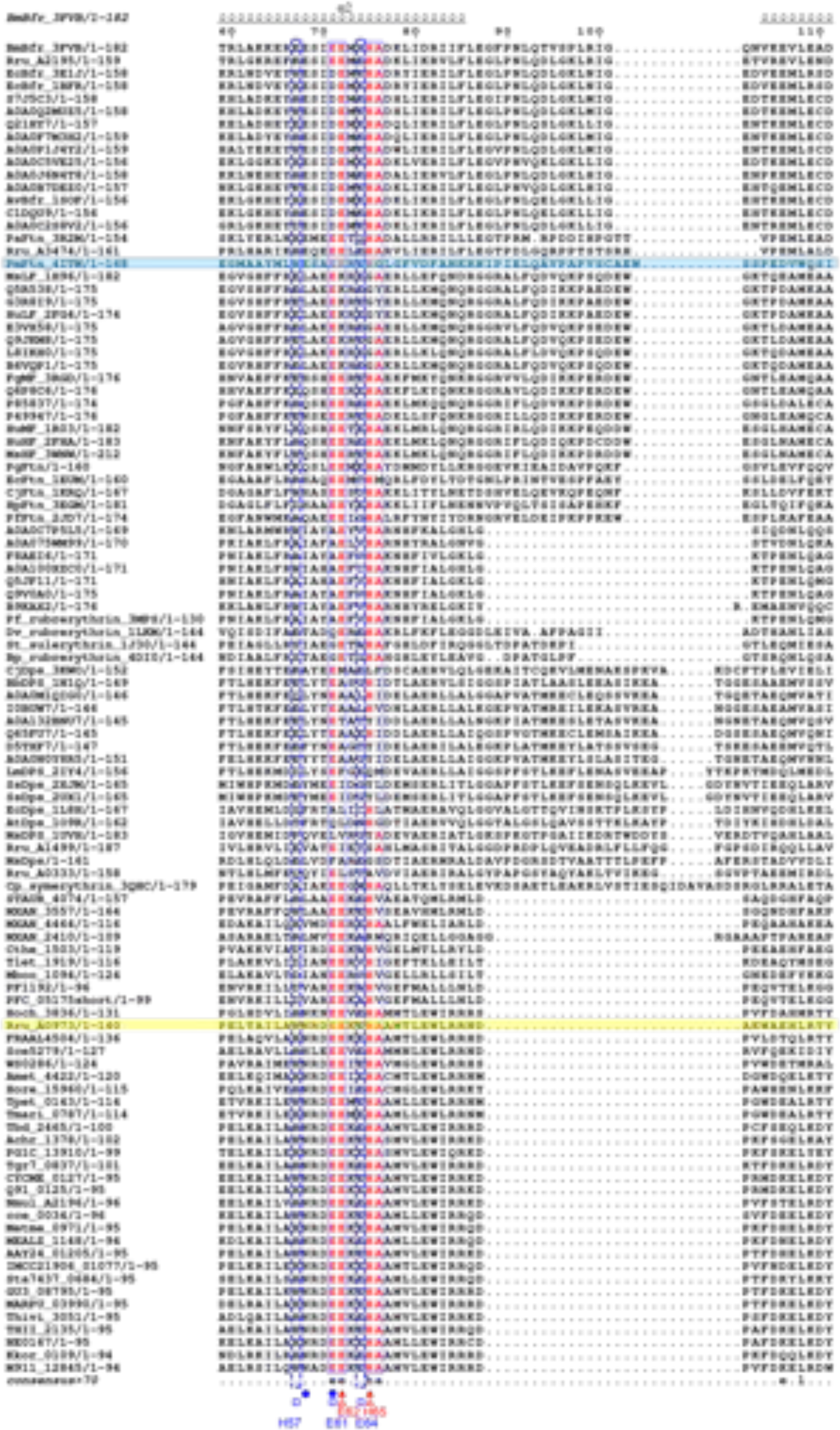

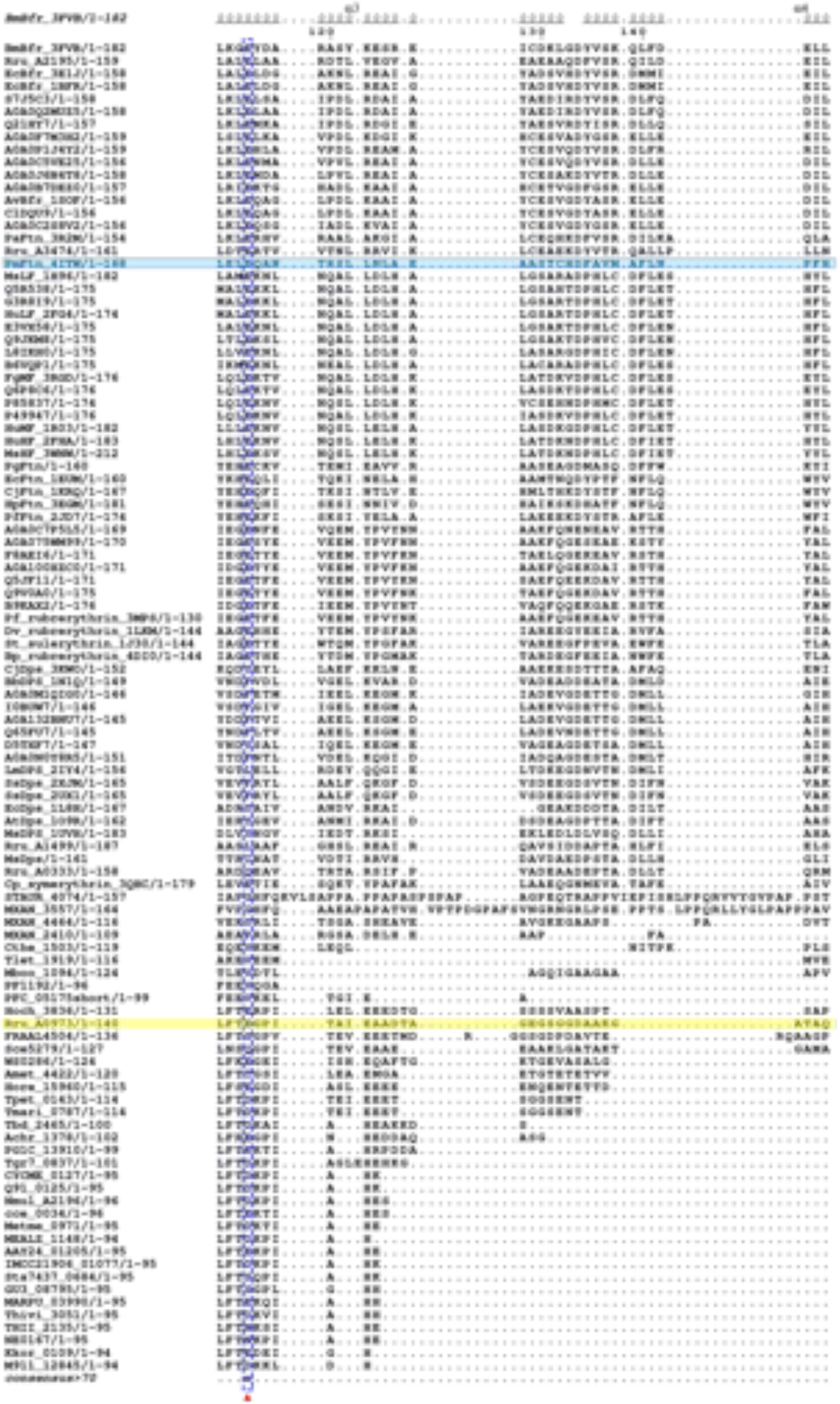

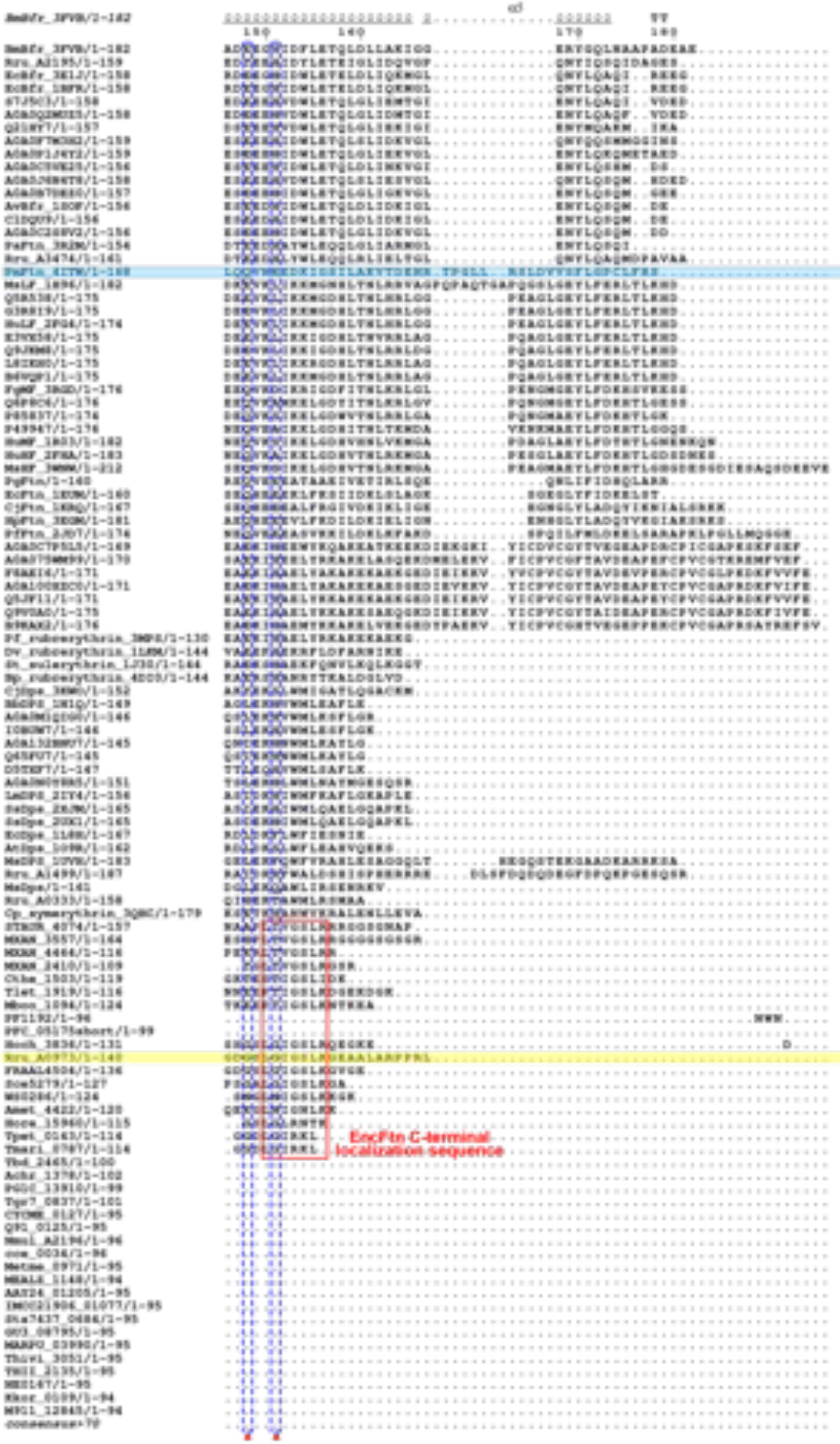

Amino acid sequences of ferritin family proteins were aligned progressively using EMBL-EBI web services^1,2^ Clustal Omega^3^, T-Coffee^4^ and MAFFT^5^. Protein names were adapted from either UniprotKB^6^, KEGG^7^ database or common name with PDB entry code^8^. Sequences were sorted in an order corresponding to the clades in phylogenetic tree (Figure 13). The alignment was edited by Espript 3.0 web server^9^. The *Rhodospirillum rubrum* EncFtn (Rru_A0973) sequence was highlighted in yellow. The ferroxidase centre (FOC) of *Pseudo-nitzschia multiseries* ferritin (PmFtn_4ITW) (highlighting in blue) consists of Fe_A_ site (E16, E49, E52) and Fe_B_ site (E49, E95, E131, Q128) which were labelled with solid red triangles^10^. Another iron binding site in PmFtn_4ITW (the gateway site or Fe_C_ site^11^) consisted of E48, E45 and E131 which are marked with solid blue circles^10^. The FOC of *R. rubrum* EncFtn was labelled with empty red triangles as E32, E62, H65 and Y39; and the iron entry site was marked with empty blue circles including E31 and E34.The putative iron exit site was marked with empty blue squares including H57, E61 and E64. The C-terminal localization sequences common to the encapsulin associated ferritins were highlighted within the red rectangle.

**Supplementary File 2.**
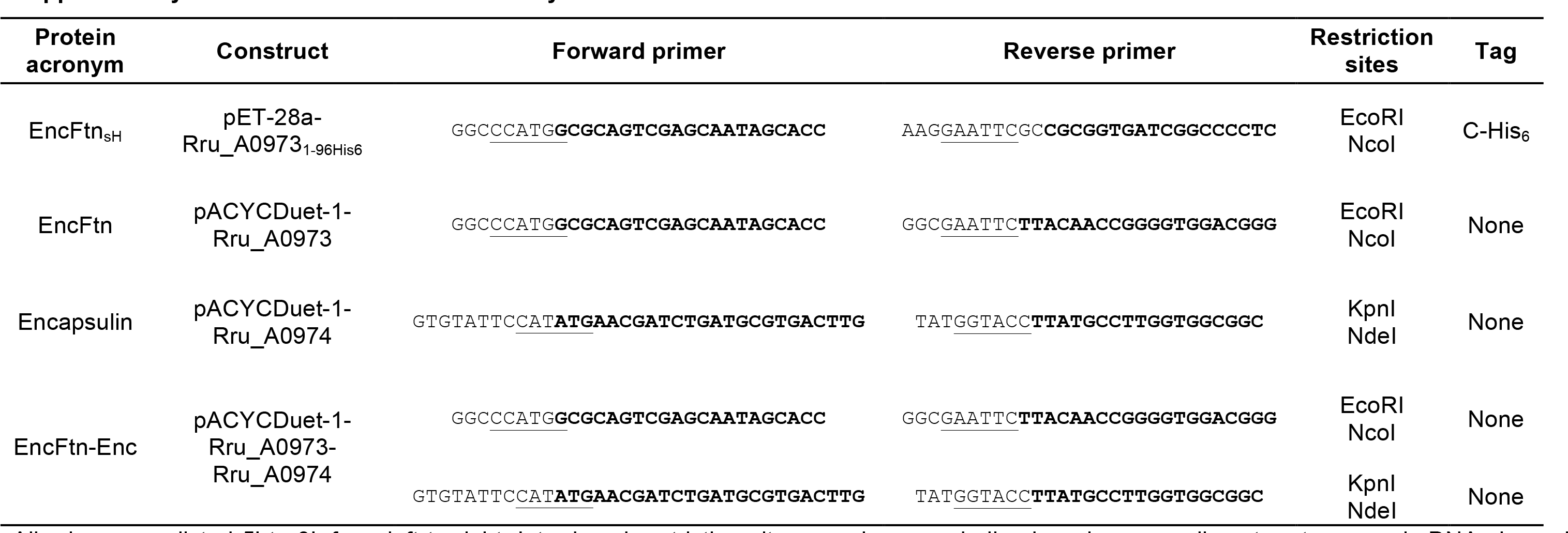
Primers used in this study.

All primers are listed 5’ to 3’, from left to right. Introduced restriction sites are shown underlined; regions complimentary to genomic DNA shown in bold.

